# Landscape Expansion Microscopy Reveals Interactions between Membrane and Phase-Separated Organelles

**DOI:** 10.64898/2025.12.10.693600

**Authors:** Yinyin Zhuang, Zhao Zhang, Zhipeng Dai, Xiaoyu Shi

## Abstract

Landscape Expansion Microscopy (land-ExM) is a light microscopy technique that visualizes both lipid and protein ultrastructural context of cells. Achieving this level of detail requires both superresolution and a high signal-to-noise ratio. Although expansion microscopy (ExM) provides superresolution, obtaining high signal-to-noise images of both proteins and lipids remains challenging. Land-ExM overcomes this limitation by using self-retention trifunctional anchors to significantly enhance protein and lipid signals in expanded samples. This improvement enables the accurate visualization of diverse membrane organelles and phase separations, as well as the three-dimensional visualization of their contact sites. As a demonstration, we revealed triple-organellar contact sites among the stress granule, the nuclear tunnel, and the nucleolus. Overall, land-ExM offers a high-contrast superresolution platform that advances our understanding of how cells spatially coordinate interactions between membrane organelles and phase separations.

**eTOC Summary:** Zhuang et al. introduce land-ExM, a super-resolution approach that simultaneously maps protein and lipid ultrastructure in cells with high contrast. This method visualizes 3D interactions between membrane-bound organelles and phase-separated condensates, uncovering organelle contact sites such as stress granules at nuclear tunnels adjacent to nucleoli.

## Introduction

Resolving the ultrastructural protein and lipid context of the cell is crucial for understanding how proteins and lipids are spatially assembled for cellular functions. It requires imaging techniques that provide both super-resolution below 100 nm and good contrast of proteins and lipids. Electron microscopy (EM) techniques, such as focused ion beam scanning electron microscopy (FIB-SEM) and cryo-electron tomography (cryoET), have provided nanometer resolution. They have therefore been the main approaches for imaging of cell context (Grunewald et al., 2003; Narayan and Subramaniam, 2015; Rigort et al., 2012; Xu et al., 2017) and membrane contact sites (MCS) (Obara et al., 2024; Wu et al., 2018). Correlative light and electron microscopy (CLEM) has further combined the strengths of electron microscopy in resolution and light microscopy in protein specificity to offer spatially resolved, multimodal insights into cellular ultrastructure (de Boer et al., 2015; Godman et al., 1960; Hauser et al., 2017). Over the past two decades, super-resolution light microscopy has rapidly narrowed the resolution gap between electron and light microscopy (Balzarotti et al., 2017; Betzig et al., 2006; Gustafsson, 2000; Hell and Wichmann, 1994; Huang et al., 2008; Pavani et al., 2009; Rust et al., 2006). More recently, expansion microscopy (ExM) has emerged as a light microscopy solution to visualize the ultrastructural protein and lipid context (Damstra et al., 2022; Karagiannis et al., 2019; Klimas et al., 2023; Mao et al., 2020; M’Saad and Bewersdorf, 2020; Shin et al., 2025; Sun et al., 2021; White et al., 2022).

By physically expanding cells or tissues, ExM allows light microscopes to achieve an effective resolution that is 3- to 20-fold higher than before expansion (Chang et al., 2017; Chen et al., 2015; Damstra et al., 2022; Ku et al., 2016; Li et al., 2022; Shaib et al., 2024; Truckenbrodt et al., 2018; Wang et al., 2024). ExM methods were initially developed for super-resolution imaging of targeted proteins (Chen et al., 2015; Chozinski et al., 2016; Ku et al., 2016; Tillberg et al., 2016) and mRNAs (Chen et al., 2016). Over time, recent advancements have introduced innovative approaches for imaging ultrastructural contexts composed of proteins, lipids, DNA, and carbohydrates (Damstra et al., 2022; Karagiannis et al., 2019; Klimas et al., 2023; M’Saad and Bewersdorf, 2020; Mao et al., 2020; Shin et al., 2025; Sun et al., 2021; White et al., 2022). These methods created EM-like images on light microscopes, which have slightly lower resolution but are more accessible than electron microscopes. For example, lipid ExM has enabled detailed imaging of lipids by integrating lipid-binding dyes into the ExM protocol (Karagiannis et al., 2019; Shin et al., 2025; White et al., 2022). Click-ExM integrates click chemistry with ExM, allowing for labeling and imaging of a wide range of biomolecules, including lipids, glycans, proteins, DNA, and RNA (Sun et al., 2021). Fluorescent Labeling of Abundant Reactive Entities (FLARE) resolved protein, carbohydrate, and DNA contexts by covalently staining cells with small fluorescent dyes followed by expansion microscopy (Mao et al., 2020). Pan-ExM (M’Saad and Bewersdorf, 2020), Ten-fold Robust Expansion Microscopy (TREx) (Damstra et al., 2022), and Magnify (Klimas et al., 2023) achieved higher expansion factors for more details of proteins, lipids, or DNA ultrastructures. Chromatin expansion microscopy (chromExM) specializes in resolving nanoscale chromatin architecture using metabolic labeling of DNA (Pownall et al., 2023). These methods have been combined with immunostaining to localize targeted proteins on contextual channels, providing an all-optical alternative to CLEM with slightly lower resolution but higher accessibility and matching resolution between specific targets and contextual structures.

Despite the progress achieved with ExM, achieving high signal-to-noise ratios simultaneously in both protein and lipid imaging remains challenging. To optimize the signal-to-noise ratios for both protein and lipid imaging, we have developed a new labeling and anchoring strategy, called landscape expansion microscopy (land-ExM). This method builds on our concept of a self-retention trifunctional label for ExM (Shi et al., 2021). The self-retention label reacts with biomolecules, anchors itself to the hydrogel, and can be fluorescently labeled after gelation. In our prior work on label retention expansion microscopy (LR-ExM), we demonstrated that antibodies conjugated with self-anchoring trifunctional labels achieved several-fold higher brightness compared with ExM using regular fluorescent antibodies (Shi et al., 2021; Zhuang and Shi, 2023). In this work, we extend this approach by employing self-anchoring probes that not only label and anchor all proteins but also crosslink lipid probes to the hydrogel. This strategy provides bright signals for both protein and lipid channels and simplifies the workflow of contextual ExM.

While the protein channel of land-ExM reveals phase separations-the membraneless compartments, the lipid channel specifies membrane organelles based on their morphologies. By leveraging both channels and combining with immunostaining, land-ExM becomes an efficient tool for identifying contact sites between membrane organelles and phase separations. In this study, we demonstrated membrane and phase separation contact sites involving three structures: the stress granule, the nuclear tunnel, and the nucleolus. This finding illustrates that land-ExM is a powerful tool for the scientific community to investigate the intricate interactions between organelles in cells and tissues.

## Results

### Principle and method development

Fig. 1A illustrates the workflow of land-ExM, which consists of eight sequential steps: (1) cell fixation, (2) staining lipids, (3) optional immunostaining, (4) anchoring proteins and lipids, and staining proteins, (5) gelation, (6) heat denaturation, (7) fluorescence labeling, and (8) expansion. The critical step in achieving a high signal-to-noise ratio for protein and lipid imaging is step (4), where the trifunctional anchor NHS-biotin-MA (Fig. 1B) simultaneously anchors proteins and the lipid probe, mCLING. NHS-biotin-MA contains three functional groups: (1) N-hydroxysuccinimide (NHS) ester, which reacts with primary amines in proteins, antibodies, and mCLING containing seven primary amines (Fig. 1C); (2) methacrylate (MA), which covalently inserts to the acrylamide polymer during gelation; and (3) biotin, which enables protein imaging by coupling with fluorescent streptavidin. Initially designed for LR-ExM (Shi et al., 2021), this trifunctional probe has been adapted for land-ExM to optimize the signal-to-noise ratio of contextual imaging.

**Fig. 1.**
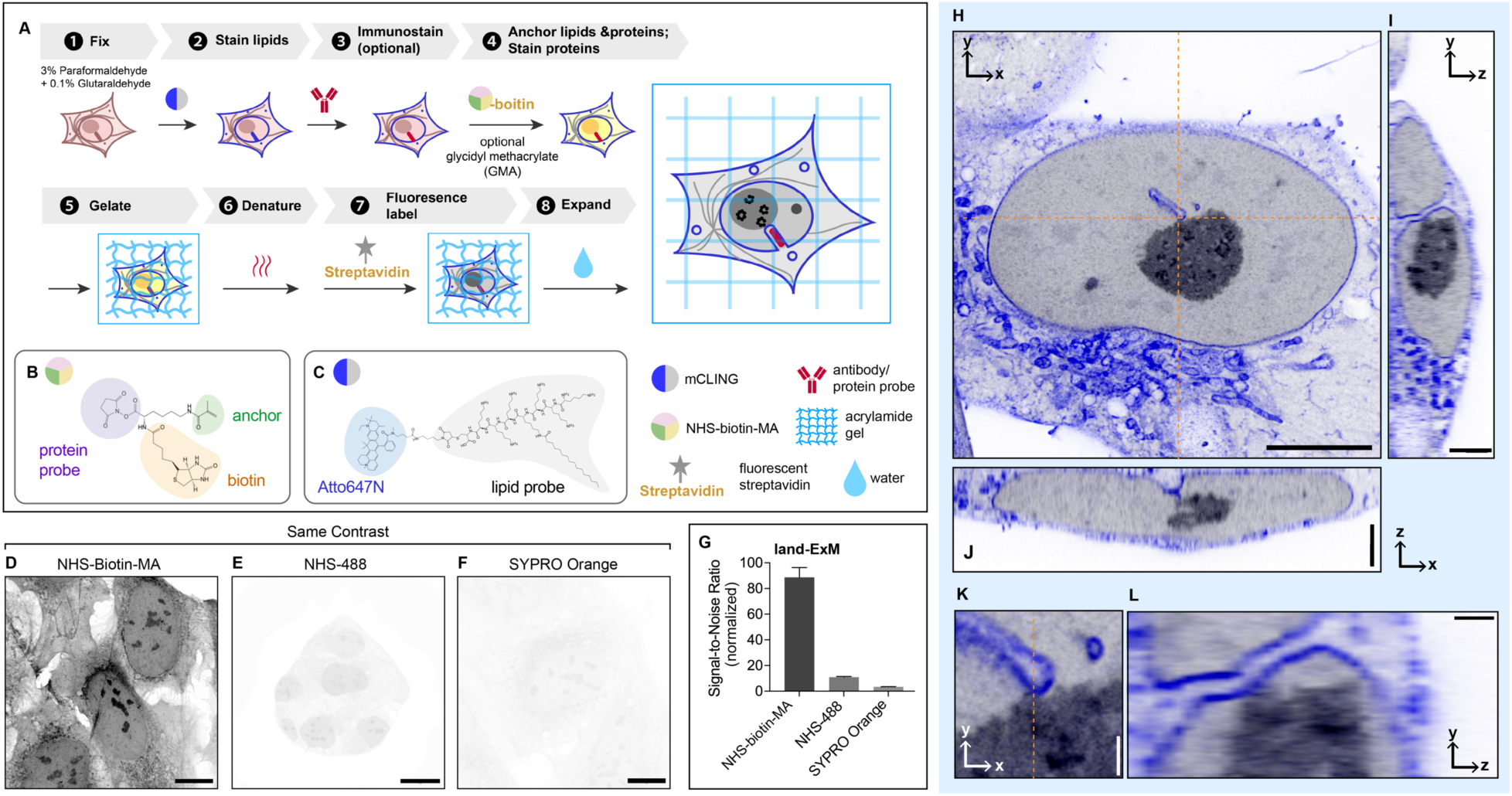
Land-ExM visualizes the protein and lipid context of cells. (A) Workflow of Land-ExM. (B) Schematic of NHS-Biotin-MA linker. (C) Schematic of mCLING. (D) Land-ExM image of U2OS cells incubated with NHS-Biotin-MA linker. Scale bar: 10 µm in pre-expansion unit. Linear expansion factor: 4. (E) ExM image of U2OS cells incubated with NHS-MA linker and stained with Alexa Fluor 488 NHS ester dye. Scale bar: 10 µm in pre-expansion unit. Linear expansion factor: 4.2. (F) ExM image of U2OS cells incubated with GMA linker and stained with SYPRO Orange. Scale bar: 10 µm in pre-expansion unit. Linear expansion factor: 4.2. (G) Bar chart comparing signal-to-noise ratios of protein context images obtained with different ExM methods shown in (D-F). The signal-to-noise ratio is calculated as the average pixel value of the area with cells divided by the average pixel value of the area without cells in each image. Each bar represents the mean ± standard error of more than 10 cells. (H-J) Different views of Land-ExM images of a breast cancer cell, UCI082014, stained with mCLING for lipid content. The orange dashed lines in (H) show where the orthogonal views (I and J) align. Scale bar: 5 µm (H), 2 µm (I and J) in pre-expansion unit. Linear expansion factor: 3.8. (K) Magnified images of (H). (L) Magnified images of (I). The orange dashed line in (K) shows where the orthogonal view (L) aligns. Scale bar: 0.5 µm in pre-expansion unit. Linear expansion factor: 3.8. All images were taken with an Airyscan microscope. Images D-F were adjusted to the same contrast.

To assess the efficacy of land-ExM in protein imaging, we compared the fluorescence signal from NHS-biotin-MA with that from standard protein dyes, including covalent NHS dyes (e.g., Alexa Fluor 488) and non-covalent stains (e.g., SYPRO Orange). The workflows for these control groups were identical to land-ExM except for steps (4) and (7), where methacrylic acid N-hydroxysuccinimide ester (NHS-MA) or glycidyl methacrylate (GMA) was used as the protein anchor in step (4), and proteins were stained with NHS-Alexa Fluor 488 or SYPRO Orange in step (7). For land-ExM, streptavidin conjugated with Alexa Fluor 488 was used to label NHS-biotin-MA in step (7). The reaction between NHS-biotin-MA and streptavidin follows a 1:1 stoichiometry, with each streptavidin molecule conjugated to an average of 0.9 dye molecules. Consequently, each NHS-biotin-MA is converted to about 1 Alexa Fluor 488 molecule in the land-ExM method, which allows for a fair comparison with the control groups.

Our results show that the land-ExM protein image has significantly higher signal-to-noise ratios than ExM using NHS-Alexa Fluor 488 and SYPRO Orange (Fig. 1D-F). The signal-to-noise ratio of the land-ExM protein image is 8 and 30 times higher than NHS-Alexa Fluor 488 and SYPRO Orange staining, respectively (Fig. 1G). It is likely because NHS-Alexa Fluor 488 (NHS-488) competes with the anchor molecule NHS-MA for primary amines, while NHS-MA consumes primary amines earlier in the workflow than NHS-488. In contrast, NHS-biotin-MA reacts with all primary amines without competition from other steps. In addition, NHS-biotin-MA anchors itself to the hydrogel, while NHS-488 relies on proteins to anchor to the hydrogel indirectly. The drawback of indirect anchoring is that NHS-488 on proteins not anchored to the hydrogel will be washed away during expansion. The self-anchoring strategy using NHS-biotin-MA avoids probe loss and further enhances the signal.

NHS-biotin-MA also anchors the lipid dye mCLING by reacting with its primary amines. Atto647N fluorescence from anchored mCLING forms the lipid channel for land-ExM. First, we optimized the mCLING staining protocols for the best labeling efficiency of lipids in cultured cells. We found that the working concentration of mCLING is important, and it needs to be optimized for each lot of the product (Fig. S1). Second, we optimized the concentration and incubation duration of NHS-biotin-MA for high-quality lipid imaging in companion with the protein channel. Fig. 1H-J depicts a 3D stack of protein and lipid images of an expanded mammalian cell with a high signal-to-noise ratio in both channels. The lipid channel (blue) reveals a nuclear tunnel, where the nuclear membrane invaginates deeply through the whole nucleus, while the protein channel (grey) highlights the nucleolus with unique phase separation. The measured lateral resolution of the Airyscan microscope is 138 nm. The 4.0 linear expansion factor of the cells used for Fig. 1 results in an effective lateral resolution of 35 nm. With this resolution and land-ExM’s high signal-to-noise ratio, contact sites between the nuclear tunnel and the nucleolus were clearly visualized (Fig. 1K&L). This observation is consistent with previous findings using electron microscopy (Bouteille and Hemon, 1979; Malhas et al., 2011). Compared with electron microscopy, land-ExM’s faster speed, 3D imaging, and multiplexity will enable a more statistical understanding of the interactions between the nuclear tunnel and the nucleolus. We explored the frequency and functions of the nuclear tunnel-nucleolus interaction with land-ExM in a recent preprint (Zhuang et al., 2024).

As NHS-biotin-MA reacts to both proteins and mCLING, there is a potential risk of cross-contamination between protein and lipid signals. To avoid crosstalk, we attempted an alternative workflow that anchors and stains proteins with NHS-Biotin-MA before the addition of mCLING (Fig. S2A). GMA is subsequently used as an additional anchor to retain the mCLING signals in gel (Fig. S2A, step 5). As expected, the NHS channel showed distinct fluorescence patterns from the mCLING channel (Fig. S2B). We compared the result of this crosstalk-free workflow (Fig. S2B) with the original workflow (Fig. S2C), finding that the risk of cross-contamination in the original workflow is negligible. It is due to the significantly lower abundance of mCLING than native proteins. Therefore, for typical cell lines and tissues, the original workflow (Fig. 1A) should be reliable. But for samples exceptionally rich in lipid, we recommend the crosstalk-free land-ExM workflow (Fig. S2A). For users focusing on lipid context but with a low requirement for protein imaging, commercial NHS-MA or glycidyl methacrylate (GMA) (Cui et al., 2023) can serve as an alternative to NHS-biotin-MA (Fig. 1A, step 4), and NHS dyes can be used for protein fluorescent labeling in step 7. This will yield an equally high signal-to-noise ratio in lipid imaging following our instructions in the Methods section, but with significantly dimmer signals in the protein channel (Fig. 1G).

### Land-ExM is compatible with proteinase K digestion

Another advantage of land-ExM is its compatibility with proteinase K digestion. Proteinase K digestion was widely used for antibody-based ExM techniques (Chen et al., 2015; Chozinski et al., 2016; Tillberg et al., 2016), which usually results in a higher expansion factor and less distortion than heat denaturation. However, most current ExM methods for protein and lipid ultrastructural imaging use heat denaturation(Damstra et al., 2022; Karagiannis et al., 2019; M’Saad and Bewersdorf, 2020; Shin et al., 2025), and are not well compatible with proteinase K digestion. It is because these methods rely on native proteins or antibodies to retain the NHS ester dye in the hydrogel. If proteinase K is used, it can fragment proteins and antibodies and cause loss of NHS ester dye on the fragments that do not crosslink to the hydrogel. However, land-ExM avoids this problem because all protein and lipid probes are covalently anchored through the NHS-biotin-MA. This advantage made land-ExM compatible with K digestion. Figure S3A showed the workflow of the proteinase K version of land-ExM. As expected, land-ExM using proteinase K retained protein and lipid signals after expansion (Fig. S3B and D). Subcellular structures identified in the heat-denaturation land-ExM workflow are identifiable using the proteinase K land-ExM workflow (Fig. S3C). The lipid signal after proteinase K digestion (Fig. S3D) is dimmer than that obtained using heat denaturation (Fig. S3E), but it can still reveal the signature morphology of most membrane organelles.

### Land-ExM enhances protein and lipid signals of TREx and pan-ExM

We applied the labeling and anchoring strategy of the land-ExM to TREx(Damstra et al., 2022) and pan-ExM(M’Saad and Bewersdorf, 2020), which are existing ExM techniques for ultrastructural protein and lipid context imaging using a higher expansion factor (7 to 20 times). By simply switching the gelation monomer to that of TREx at step 5 in Fig. 1A, we achieved an expansion factor of 7.0. We term this land-ExM variant land-TREx (Fig. 2B and E). Compared with TREx (Fig. 2C and F), land-TREx showed a fivefold increase in signal-to-noise ratio in the protein channel and a twofold increase in the lipid channel (Fig. 2D and G). This improvement accounts for the use of NHS-biotin-MA. To integrate land-ExM with pan-ExM (land-pan-ExM), we introduced NHS-Biotin-MA right after cell fixation (Fig. 2A). In the original pan-ExM workflow, the acrylamide and formaldehyde incubation step was designed to anchor proteins to the hydrogel and to prevent inter-protein crosslinking. This step consumes primary amines of proteins, which competes with the reaction between NHS ester dye and primary amine at a later step and thus hinders the protein labeling efficiency. However, land-pan-ExM avoids this competition by labeling and anchoring the proteins through NHS-biotin-MA beforehand. As a result, the signal-to-noise ratio of the protein channel of land-pan-ExM was 2 times higher than that of pan-ExM (Fig. 2H to J). In addition, land-pan-ExM showed a threefold increase in lipid signal compared with pan-ExM. (Fig. 2K to M). This is because NHS-Biotin-MA anchors mCLING to the hydrogel in addition to formaldehyde. Overall, we showed that land-ExM is compatible with other ExM techniques and enhances protein and lipid signals.

**Fig. 2.**
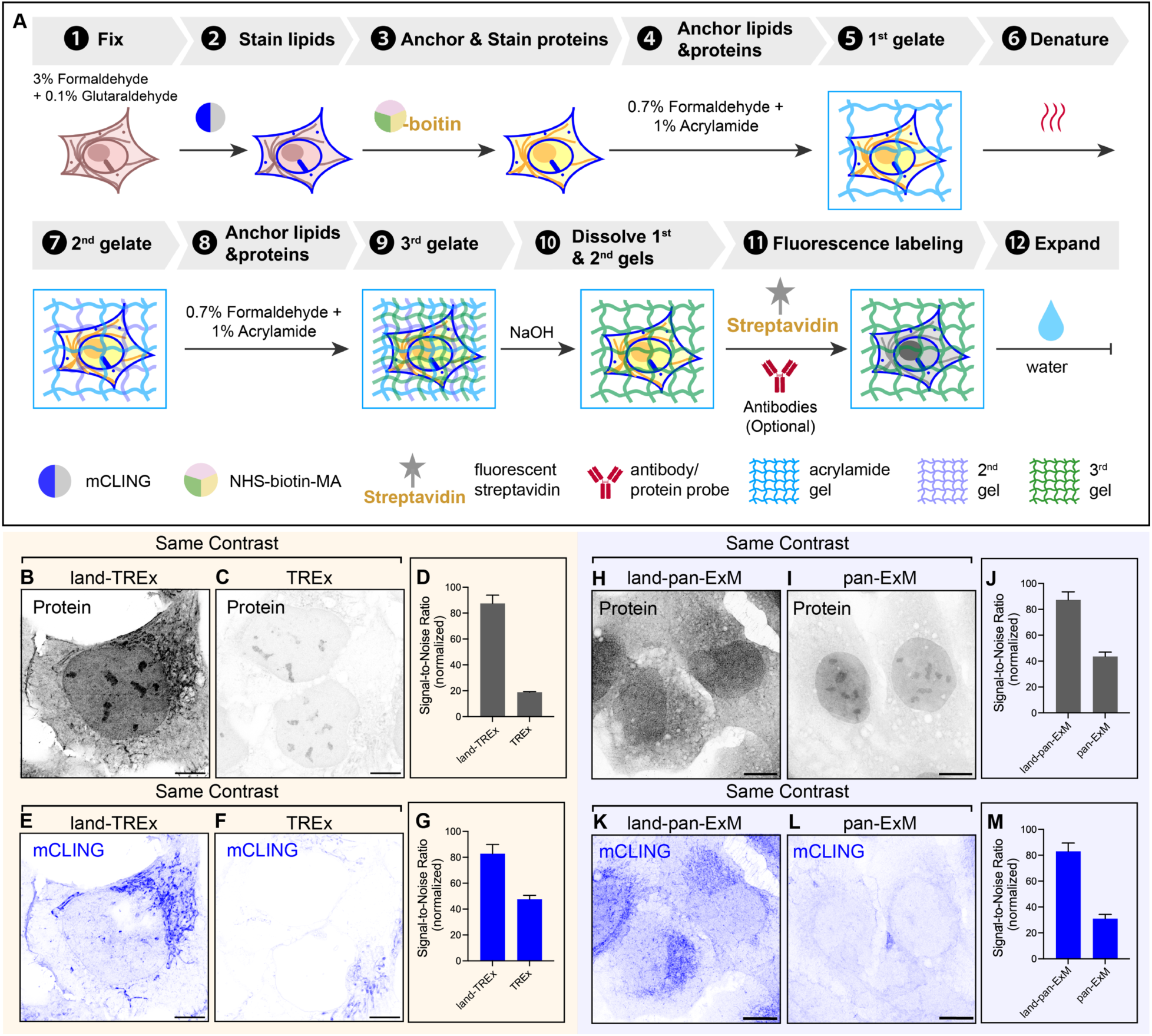
Land-ExM labeling and anchoring strategies improve the signal of TREx and pan-ExM. (A) Workflow of land-pan-ExM, which only replaces the labeling strategy of pan-ExM with the labeling strategy of land-ExM. (B) land-TREx protein channel of U2OS cells, where proteins were labeled and anchored with NHS-Biotin-MA. Scale bar: 5 µm in pre-expansion unit. Linear expansion factor: 7. (C) TREx protein channel of U2OS cells, where proteins were anchored with acryloyl-X SE and stained with Alexa Fluor 488 NHS ester. Scale bar: 5 µm in pre-expansion unit. Linear expansion factor: 7. (D) Bar chart comparing the signal-to-noise ratio of the protein channel in land-TREx and TREx. The signal-to-noise ratio is calculated as the average pixel value of the area with cells divided by the average pixel value of the area without cells in each image. Each bar represents the mean ± standard error of more than 20 cells. (E) land-TREx lipid channel of U2OS cells, where lipids were labeled by mCLING and anchored with NHS-Biotin-MA. Scale bar: 5 µm in pre-expansion unit. Linear expansion factor: 7.0. (F) TREx lipid channel of U2OS cells, where lipids were anchored with acryloyl-X SE and stained with mCLING. Scale bar: 5 µm in pre-expansion unit. Linear expansion factor: 7.0. (G) Bar chart comparing the signal-to-noise ratio of the lipid channel of land-TREx and TREx. The signal-to-noise ratio is calculated as the average pixel value of the area with cells divided by the average pixel value of the area without cells in each image. Each bar represents the mean ± standard error of more than 20 cells. (H) land-pan-ExM protein channel of U2OS cells, where proteins were labeled and anchored with NHS-Biotin-MA. Scale bar: 5 µm in pre-expansion unit. Linear expansion factor: 12.0. (I) pan-ExM protein channel of U2OS cells labeled with Alexa Fluor 488 NHS ester. Scale bar: 5 µm in pre-expansion unit. Linear expansion factor: 12.0. (J) Bar chart comparing the signal-to-noise ratio of the protein channel in land-pan-ExM and pan-ExM. The signal-to-noise ratio is calculated as the average pixel value of the area with cells divided by the average pixel value of the area without cells in each image. Each bar represents the mean ± standard error of more than 20 cells. (K) land-pan-ExM lipid channel of U2OS cells, where lipids were stained following the workflow (A). Scale bar: 5 µm in pre-expansion unit. Linear expansion factor: 12.0. (L) pan-ExM lipid channel of U2OS cells labeled with mCLING. Scale bar: 5 µm in pre-expansion unit. Linear expansion factor: 12.0. (M) Bar chart comparing the signal-to-noise ratio of the lipid (mCLING) channel in land-pan-ExM and pan-ExM. The signal-to-noise ratio is calculated as the average pixel value of the area with cells divided by the average pixel value of the area without cells in each image. Each bar represents the mean ± standard error of more than 20 cells. All images were taken with an Airyscan microscope.

### Land-ExM visualizes membrane organelles and phase separation

Land-ExM can visualize membrane structures and phase-separated structures based on their morphologies and locations in both protein and lipid channels. Here, we define phase separation as membraneless protein condensates. Fig. 3A-G display a gallery of land-ExM protein images of phase-separated compartments, such as nucleoli (Fig. 3A), nuclear bodies (Fig. 3B), and stress granules (Fig. 3C). Within the nucleus, chromatin (Fig. 3D) and nuclear pore complexes (Fig. 3E) were resolved, while in the cytoplasm, mitochondria (Fig. 3F) and cytoskeleton (Fig. 3G) were visualized. A contributor to the cytoskeleton structure in the protein channel is actin, which was confirmed with phalloidin staining (Fig. S6). Similarly, lipid land-ExM imaging distinguishes membrane structures (Fig. 3H-P), such as lipid vesicles (Fig. 3I), mitochondria (Fig. 3J), filopodia (Fig. 3K), nuclear membrane invaginations (Fig. 3L), and Golgi apparatuses (Fig. 3M). Even the trans and cis faces of a Golgi apparatus can be discernible based on lipid curvature (Fig. 3M).

**Fig. 3.**
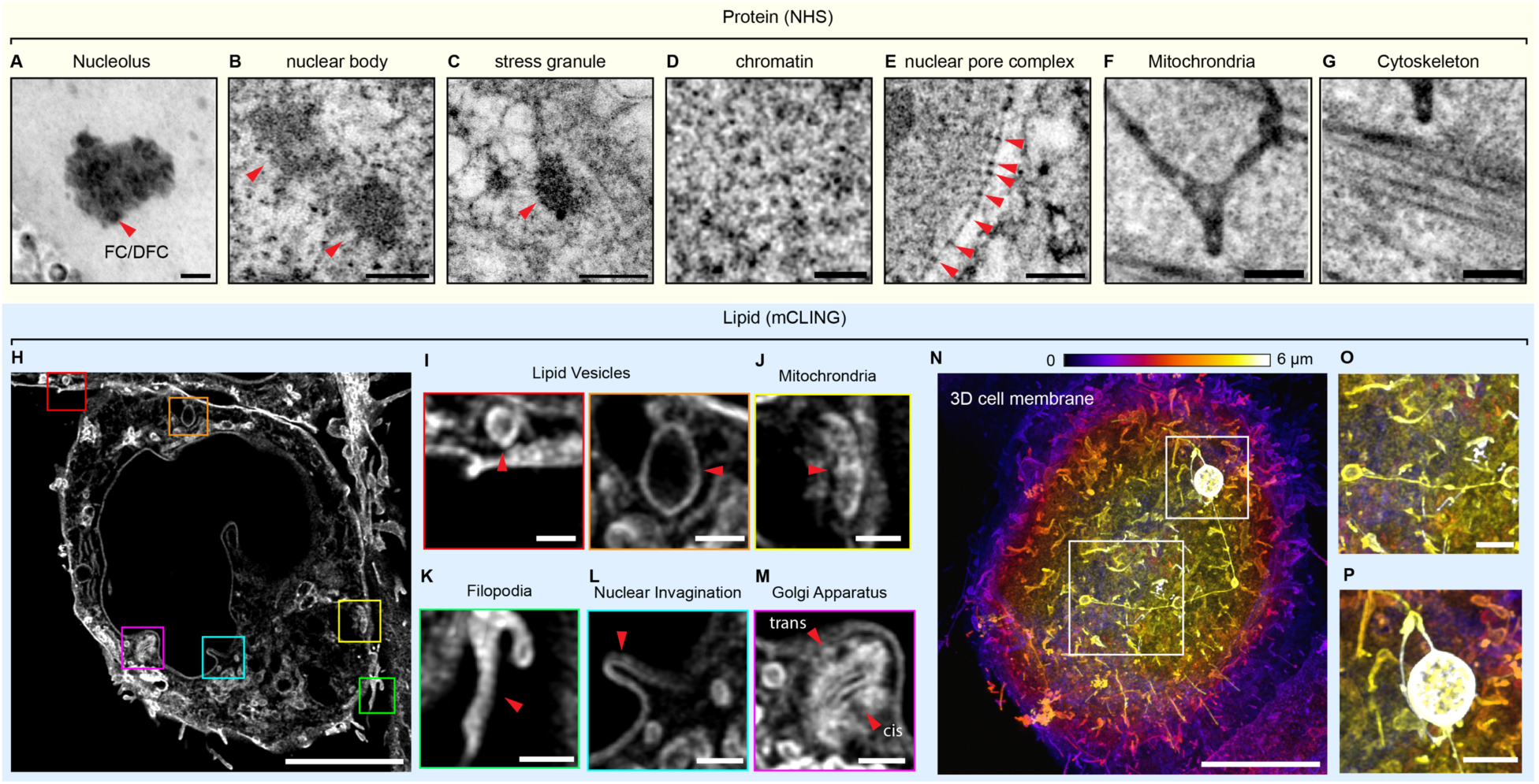
Land-ExM visualizes phase-separated and membrane organelles. (A-G) land-ExM protein images of membraneless phase separation structures. The proteins were labeled with NHS-biotin-MS, and post-gelation stained with streptavidin-Alexa Fluor 488. (A) land-ExM protein image of nucleoli in a U2OS cell. Red arrowheads indicate the fibrillar center (FC) or dense fibrillar component (DFC) of the nucleolus. Scale bar: 1 µm in pre-expansion unit. Linear expansion factor: 4.0. (B) land-ExM protein image of nuclear bodies of breast cancer cell, UCI082014. Red arrowheads indicate the nuclear bodies. Scale bar: 1 µm in pre-expansion unit. Linear expansion factor: 4.2. (C) land-ExM protein image of stress granules of a U2OS cell treated with NaAsO_2_ for 20 minutes. The red arrowhead indicates a stress granule. Scale bar: 1 µm in pre-expansion unit. Linear expansion factor: 4.0. (D) land-ExM protein image of chromatin of a breast cancer cell. Scale bar: 500 nm in pre-expansion unit. Linear expansion factor: 4.2. (E) land-ExM protein image of nuclear pore complexes of a breast cancer cell. Scale bar: 1 µm in pre-expansion unit. Linear expansion factor: 4.2. (F and G) land-ExM protein images of mitochondria and cytoskeleton of a U2OS cell. Scale bar: 1 µm in pre-expansion unit. Linear expansion factor: 4.0. (H-P) land-ExM lipid images of membrane structures. The lipids were labeled with mCLING-Atto647N. (H) land-ExM lipid image of breast cancer cell. Scale bar: 5 µm in pre-expansion unit. Linear expansion factor: 4.0. (I-M) magnified images of (H) showing different membrane structures: lipid vesicles (I), mitochondria (J), filopodia (K), nuclear invagination (L), and Golgi apparatus (M). Scale bar: 1 µm (I-M) in pre-expansion unit. (N) 3D land-ExM lipid image of a breast cancer cell after maximum intensity projection, showing the cell membrane. Color-coded by the z-dimension slices from bottom to top. Color bar: purple to white: 0 to 6 µm in pre-expansion unit. Scale bar: 5 µm in pre-expansion unit. Linear expansion factor: 4.0. (O and P) magnified images of (N) showing detailed structures of the cell membrane. Scale bar: 1 µm in pre-expansion unit. All images were taken with an Airyscan microscope.

In addition to the protein and lipid channels, immunostaining can be added to specify the organelles further (Fig. S4, S5 and S6). For example, using anti-Lamp2 and anti-clathrin antibodies, lysosomes and clathrin-coated pits can stand out from other lipid vesicles (Fig. S4). The 3D land-ExM lipid image can reveal the whole cell membrane with intricate structures (Fig. 3N), such as filopodia, cytonemes (Fig. 3O), and potential exosomes (Fig. 3P). With bright signals in protein, lipid, and immunostaining, land-ExM is optimized for discovering interactions between specific membrane organelles and phase separation.

### Triple-organellar contact sites among the stress granule, the nuclear tunnel, and the nucleolus

Studying organellar contact sites is important because these regions are not just passive points of membrane proximity, but are active hubs for communication, coordination, and metabolic regulation inside cells. The region between two organelles that come into proximity, typically separated by only 10-80 nanometers, is considered a contact site (Raimondi et al., 2023; Scorrano et al., 2019). Multi-color 3D land-ExM with lipid, protein, and antibody channels offers the specificity and resolution needed to identify organellar contact sites between organelles, including both membrane and membraneless organelles. We measured the distance between the membranes of two organelles or between a membrane and the edge of a phase-separated organelle as a metric to discriminate a contact site.

As a demonstration, we investigated stress granules (SGs), which suppress mRNA translation by sequestering mRNAs into phase-separated compartments in the cytoplasm. To form stress granules, we treated the cells in 500 µM sodium arsenite for 20 and 60 minutes, respectively, and compared them to the ones without treatment (Fig. S7). We immunostained the cells with stress granule marker G3BP1 and visualized it in the context of proteins and lipids using the multi-color 3D land-ExM. When the cells were not treated with sodium arsenite, G3BP1 diffused across the cell (Fig. S7A and D). When the cells were incubated in a sodium arsenite solution, G3BP1 condensates appeared (Fig. S7B, C, E, and F). And when we removed the sodium arsenite, the G3BP1 condensates disappeared. These results validate that the G3BP1 condensates are caused by stress, and G3BP1 can be used as a stress granule marker, which is consistent with how sodium arsenite was used in previous research to introduce stress granules (Wheeler et al., 2016). Surprisingly, some SGs appeared near the nucleolus (boxed in Fig. 4A and magnified in Fig. 4B-E) instead of cytoplasm (Fig. S7G). We asked whether these two phase-separated organelles directly interact in the nucleus. The answer is no when considering the lipid context. The lipid channel showed that these small SGs were localized inside nuclear tunnels, separated from the nucleolus by the nuclear membrane (Fig. 4F-K). The nuclear tunnel forms two contact sites on its cytoplasmic side and nucleoplasmic side (Fig. 4G&H). It contacts the SG on the cytoplasmic side and with the nucleolus on the nucleoplasmic side. This spatial relationship was commonly seen in nuclear tunnels of cells under stress in our experiments. We examined 114 nuclear tunnels in more than 20 cells, finding 83% tunnels contain SGs (Fig. 4R). Among the tunnels containing SGs, 60% contact nucleoli (Fig. 4S). The triple-organelle interactions may result in efficient reduction of mRNAs as a response to stress. We will discuss the potential mechanisms in the discussion section. Furthermore, we quantitatively examined whether the sodium arsenite treatment alters the nuclear tunnels. We found that U2OS cells can have 1 to 10 nuclear tunnels per nucleus, with an average number of 5 tunnels per nucleus (Fig. S7H). The diameter of the nuclear tunnels ranges from 140 nm to 400 nm, with an average of around 250 nm (Fig. S7I). Sodium arsenite treatment had no significant effect on the number or size of nuclear tunnels (Fig. S7H and I).

**Fig. 4.**
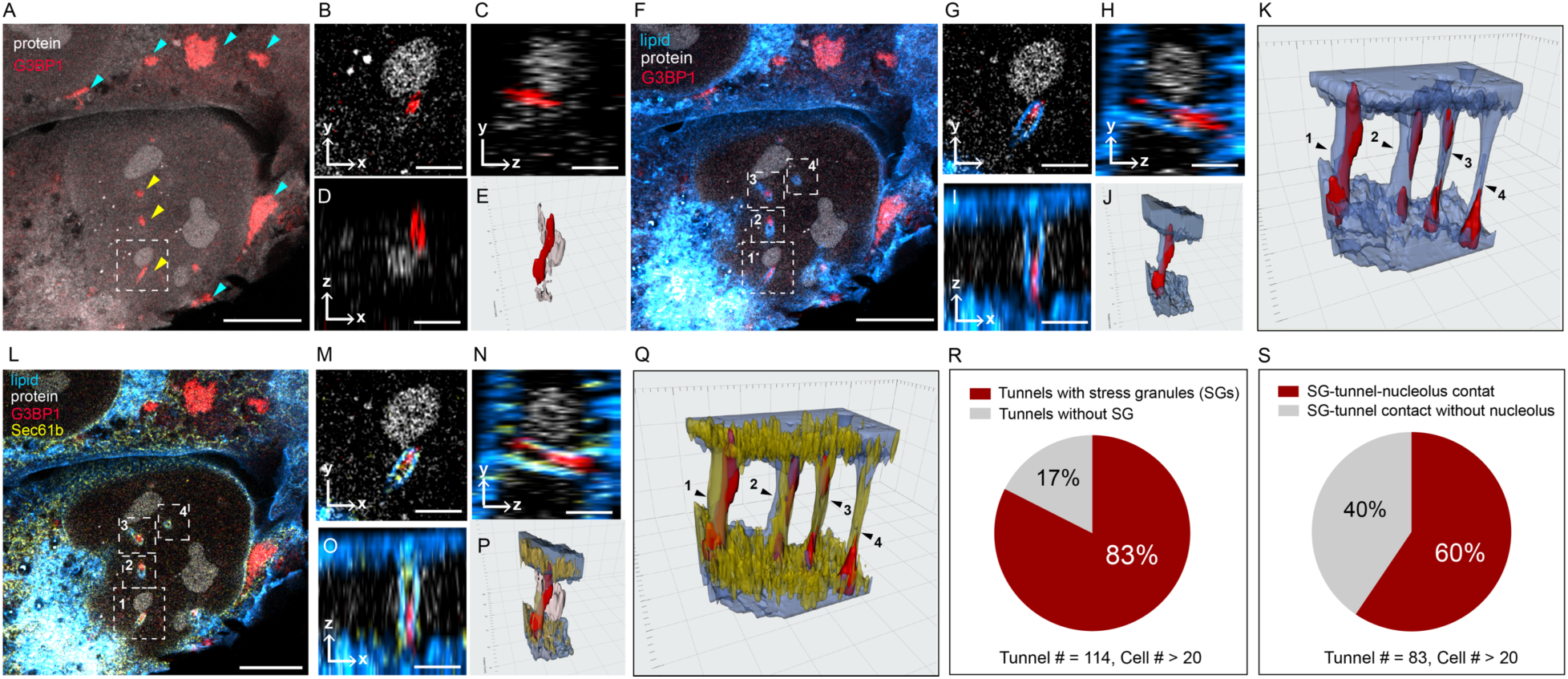
The nuclear tunnel forms a triple-organellar contact site that includes the stress granule, the nucleolus, and itself. (A) Land-ExM protein (grey) image of U2OS cells immunostained with anti-G3BP1 (red) antibody. Cells were treated with NaAsO_2_ for 1 hour. Scale bar: 5 µm in pre-expansion unit. Linear expansion factor: 4. (B-D) Different views of stress granule in the white dashed box of (A). Scale bar: 1 µm in pre-expansion unit. (E) 3D rendering of stress granule in the white dashed box of (A). In the reference grid, the spacing of major and minor tick marks is 0.5 µm and 0.1 µm in pre-expansion unit. (F) Land-ExM protein (grey) and lipid (blue) image of U2OS cells immunostained with anti-G3BP1 (red) antibody. Cells were treated with NaAsO_2_ for 1 hour. Scale bar: 5 µm in pre-expansion unit. Linear expansion factor: 4.(G-I) Different views of stress granule in the white dashed box 1 of (F). Scale bar: 1 µm in pre-expansion unit. (J) 3D rendering of stress granule in the white dashed box 1 of (F). In the reference grid, the spacing of major and minor tick marks is 0.5 µm and 0.1 µm in the pre-expansion unit. (K) 3D rendering of stress granules in the white dashed box 1-4 of (F). In the reference grid, the spacing of major and minor tick marks is 0.5 µm and 0.1 µm in the pre-expansion unit. (L) Land-ExM protein (grey) image of U2OS cells immunostained with anti-G3BP1 (red) and anti-Sec61b (yellow) antibodies. Cells were treated with NaAsO_2_ for 1 hour. Scale bar: 5 µm in pre-expansion unit. Linear expansion factor: 4. (M-O) Different views of stress granule in the white dashed box 1 of (L). Scale bar: 1 µm in pre-expansion unit. (P) 3D rendering of stress granule in the white dashed box 1 of (L). In the reference grid, the spacing of major and minor tick marks is 0.5 µm and 0.1 µm in pre-expansion unit. (Q) 3D rendering of stress granules in the white dashed box 1-4 of (L). In the reference grid, the spacing of major and minor tick marks is 0.5 µm and 0.1 µm in pre-expansion unit. (R) Pie chart of nuclear tunnels with or without SGs. Total tunnels analyzed: 114. (S) Pie chart of SG-filled nuclear tunnels that contact nucleoli versus those that do not. Total tunnel analyzed: 83. All images were taken with an Airyscan microscope.

We also observed that the endoplasmic reticulum (ER), as a part of the nuclear membrane of nuclear tunnels, was adjacent to the SGs. In our four-color land-ExM images, which captured lipids, proteins, and immunostained SG and ER markers (Fig. 4L), the ER was found adjacent to SGs (Fig. 4M-P). ER-SG contacts inside nuclear tunnels were frequently observed in cells under stress in our experiments. All four nuclear channels shown in Fig. 4Q displayed contact sites between SGs and the ER. Previous studies reported the contact between SGs and ER in the cytosol, which contributes to the SG’s formation (Liu et al., 2024; Nicchitta, 2024; Pincus and Oakes, 2024) and the ER’s response to stress (Liu et al., 2024). Our observations in nuclear tunnels highlight parallels between the SG-ER interaction within nuclear tunnels and in the cytoplasm. Our land-ExM imaging shows that the nuclear tunnels provide confined space that enables contact between the membrane and phase-separated organelles, including the ER, SGs, and nucleoli. Our recent study demonstrated that nuclear tunnels regulate ribosome biogenesis by interacting with the nucleolus (Zhuang et al., 2024). The triple contact among the SG, the nuclear tunnel, and the nucleolus may indicate more roles of nuclear tunnels under stress.

## Discussion

### Advantages of land-ExM

In this study, we developed a new expansion microscopy approach, land-ExM, that significantly enhances the signal-to-noise ratio in imaging protein and lipid ultrastructure. A key innovation in our method is the use of the trifunctional anchor NHS-biotin-MA, which simultaneously anchors proteins and the lipid probe mCLING to the hydrogel. This self-anchoring strategy results in signal-to-noise ratios significantly higher than those achieved with conventional protein dyes such as NHS-Alexa Fluor 488 or SYPRO Orange. The self-anchoring strategy is compatible with both heat denaturation and proteinase K digestion, enabling the integration of land-ExM with most ExM techniques. In this work, we demonstrated on the standard x4 ExM, TREx, and pan-ExM.

Another strength of land-ExM is its compatibility with targeted protein imaging techniques. These include immunostaining, fluorescent proteins, and self-labeling tags. Multicolor land-ExM imaging of targeted molecules, lipids, and proteins enables the precise localization of molecules within the 3D architectures formed by interacting organelles and protein complexes. The combination of contextual and targeted imaging is streamlined in land-ExM, as shown in Fig. 1A. All imaging channels are captured on the same microscope, ensuring similar resolution and straightforward alignment of all channels.

Compared with correlative light and electron microscopy (CLEM), land-ExM as well as other contextual ExM methods are more accessible, cost-effective, and faster. Its affordability and ease of use make it an attractive option for laboratories with limited resources.

Collectively, these advances in anchoring efficiency, signal brightness, multi-color imaging, and super-resolution capability position Land-ExM as a powerful method for the detailed investigation of protein and lipid ultrastructure in biological systems.

### Limitations of Land-ExM and solutions

Despite its significant advantages, land-ExM has several limitations. While its resolution surpasses that of conventional light microscopy, it is still lower than electron microscopy techniques such as cryoET, FIB-SEM, and transmission electron microscopy (TEM). This limitation hinders the method’s ability to resolve fine structural details below 10 nm, such as distinguishing between the inner and outer nuclear membranes. It is technically possible to push the resolution of land-ExM slightly beyond 10 nm by imaging with single-molecule localization microscopes, like STORM (Shi et al., 2021) and MINFLUX (Balzarotti et al., 2017; Schmidt et al., 2021), or employing more swellable hydrogel (Chang et al., 2017),(Wang et al., 2024),(Shaib et al., 2024). However, the ultimate resolution that ExM can achieve is limited by the pore size of the hydrogel, and the nanoscale distortion caused by expansion must be carefully examined.

Land-ExM is not a label-free technique, and so the contextual information it provides can be influenced by the choice of labeling probes. For protein staining, NHS ester probes give a stronger signal to lysine-rich proteins, potentially introducing bias in the representation of cellular components. To address this, alternative labeling chemistries can be employed. For example, SYPRO Orange, which interacts with the hydrophobic regions of proteins exposed after denaturation, offers complementary insights. Similarly, for lipid staining, different probes exhibit varying labeling efficiency. mCLING is well retained by membranes after fixation and permeabilization, while styryl (FM) dyes are largely lost (Revelo et al., 2014). It can therefore be used in combination with immunostaining, meeting the needs of ExM. Probes specifically developed for ExM, such as those from the Boyden lab, offer improved fluorescence retention for lipid imaging (Shin et al., 2025).

Another challenge is the potential for fixation artifacts, as land-ExM relies on chemical fixation. Fixation conditions optimized for one biomolecule may compromise the labeling efficiency of another. For example, lipid staining often requires glutaraldehyde fixation, which can mask certain protein epitopes and inhibit their immunostaining. These limitations restrict the range of experiments that can be performed using land-ExM. In addition, harsh chemical fixation can cause distortion of cellular ultrastructure at the sub-micrometer scale. One promising alternative, highlighted in recent protocols(Laporte et al., 2022), is cryofixation, which more effectively preserves native cellular structures. To assess the subcellular distortion in expanded samples, quantitative measurement of sub-micrometer distortion should be acquired for ExM experiments, particularly when using a new protocol or expanding a different organelle. GelMAP(Damstra et al., 2023), along with methods that use fiducial markers(Bianchini et al., 2021; Scheckenbach et al., 2020; Vanheusden et al., 2020), offers a solution for measuring local distortions in 2D. However, a technical gap still exists for the expansion factor mapping of whole cells in 3D.

### The nuclear tunnel organizes membrane-phase separation contact

We recently reported that nuclear tunnels can activate the nucleoli they contact, leading to increased ribosomal subunit production in nucleoli (Zhuang et al., 2024). These subunits are then exported through nuclear pore complexes (NPCs) located on the nuclear tunnels, thereby elevating mRNA translation in the cell. In this study, we observed SGs adjacent to nucleoli, separated by these nuclear tunnels, in cells under stress. As SGs primarily function to silence mRNA translation by sequestering mRNAs within their phase-separated environments, the close proximity of SGs to nucleoli via nuclear tunnels suggests a functional link. Our hypothesis is that by localizing SGs in the nuclear tunnels, the cell creates a regulatory microenvironment that can capture freshly exported mRNAs and ribosomal subunits—thereby reducing translation. This arrangement may enable cells to efficiently coordinate ribosome production, mRNA sequestration, and protein synthesis in response to stress.

While G3BP1’s classic role involves SG assembly, it also contributes to membrane repair (King et al., 2025). For example, G3BP1 is reported to repair lysosomes through a “plugging” mechanism (Bussi et al., 2023). This repair activity positions G3BP1 as a crucial integrator of cellular stress responses across both membraneless and membrane-bound compartments. It is possible that the localization of G3BP1 in the nuclear tunnels is related to these prior studies.

## Conclusion

Land-ExM represents a significant advancement in super-resolution microscopy by combining contextual ExM imaging with a self-retention trifunctional labeling strategy. This approach enables high signal-to-noise detection of both proteins and lipids while being compatible with immunostaining. In turn, land-ExM provides detailed insights into complex spatial relationships between membrane-bound and phase-separated structures. Notably, it has uncovered contact sites involving three structures: the stress granule, the nuclear tunnel, and the nucleolus. These results offer new perspectives on organelle-organelle interactions.

With its superresolution, compatibility with existing ExM protocols, and greater accessibility relative to electron microscopy, land-ExM is a practical yet powerful solution for investigating the molecular landscape of cells. By enabling researchers to visualize the protein and lipid context in cells and tissues on light microscopes, land-ExM can accelerate discoveries in organellar interactions and structure-function relationships. Future efforts to enhance its resolution, fixation methods, and labeling approaches will likely further expand its impact and applicability.

## Methods

### Key Resources Table

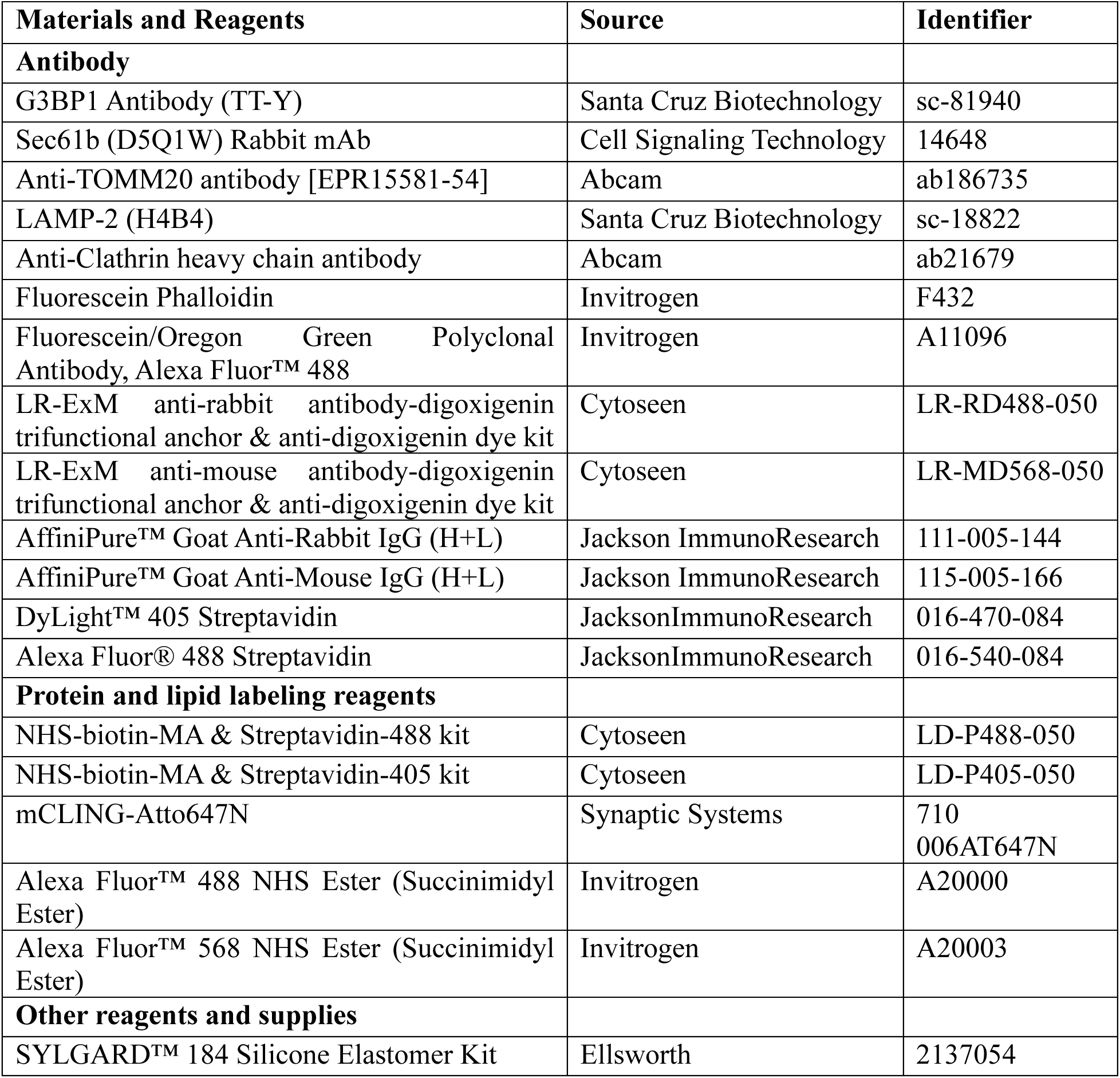
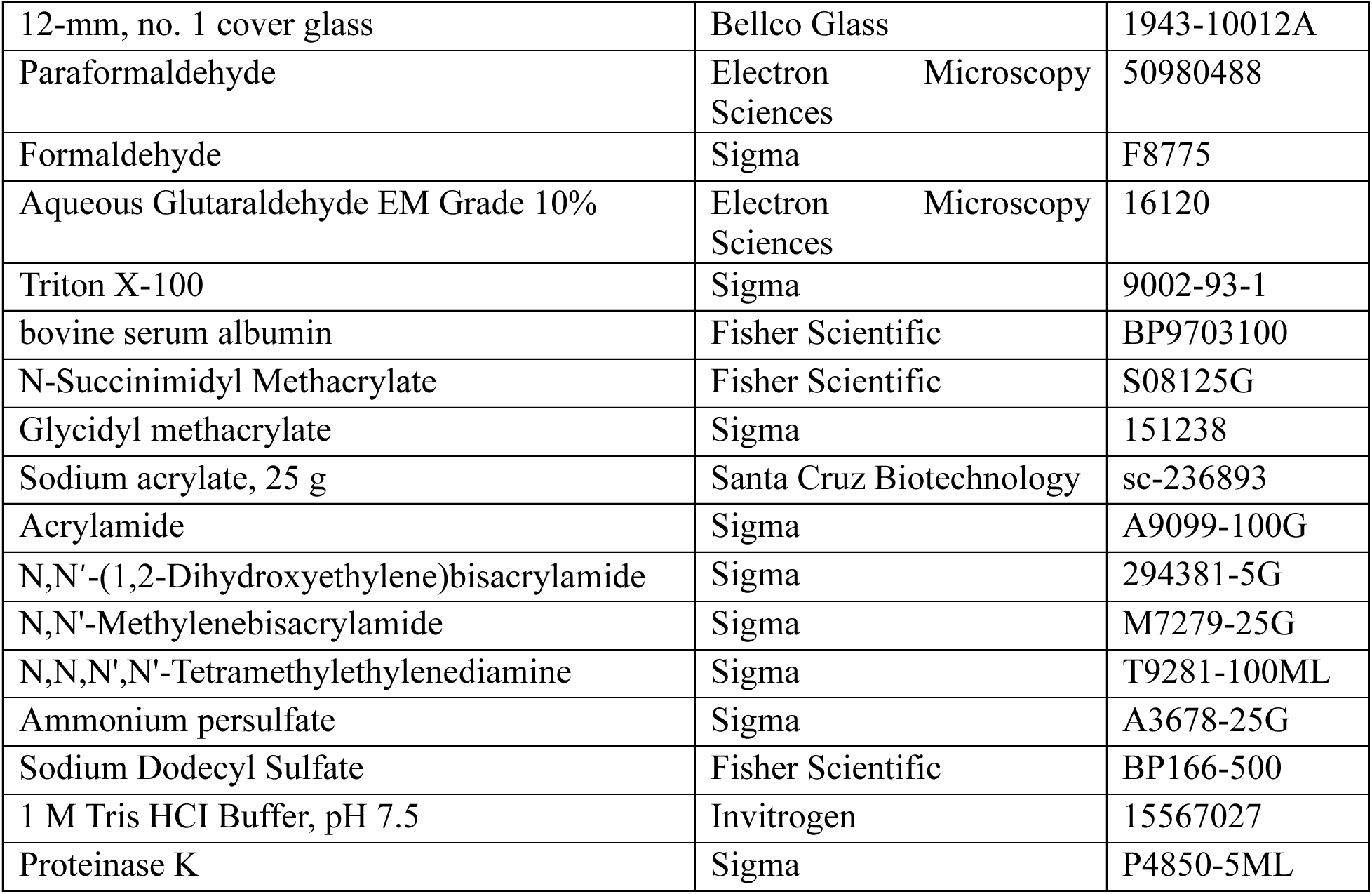

### Cell culture and Treatment

All cell lines were maintained in humidified incubator with 5% CO_2_ at 37 °C. U2OS cells were cultured in Mccoy’s 5A medium (Gibco^TM^, Cat#16600082) supplemented with 10% Fetal Bovine Serum (Gibco^TM^, cat#10082147) and 1% penicillin-streptomycin-amphotericin B (Sigma, cat#A5955). UCI082014 breast cancer cells were cultured in DMEM-high Glucose supplemented with GlutaMAX^TM^, 10% Fetal Bovine Serum and 1% penicillin-streptomycin-amphotericin B. All cell lines were tested mycoplasma-free using MycoStrip™-Mycoplasma Detection Kit (InvivoGen, cat# rep-mysnc-50) and used at less than 10 passages from thaw. UCI082014 cell was a gift from Dr. Olga V Razorenova at University of California, Irvine.

To induce stress granules in the U2OS cells, 500 µM sodium arsenite (Fisher Scientific, cat#AA41533AP) was added to the culture medium, and incubated with cells at 37 °C for 20 min to 1 hour. Cells were quickly fixed after the sodium arsenite treatment.

### Optimization of lipid staining with mCLING

We found that the mCLING-atto647N purchased may have batch-to-batch variations in purity or dye-to-mCLING ratio, which resulted in different lipid labeling efficiencies (Figure S1). The powder of mCLING-atto647N should be blue in color. Be aware that if the powder is received in white, such mCLING has limited lipid staining capability (Fig. S1A). Optimization of the dilution factor should be performed on cells before applying it to the land-ExM procedure. At low concentrations, mCLING-atto647N can only stain the plasma membrane of the cells (Figure S1A & B). However, at high concentrations, mCLING-atto647N can stain the inner membrane structures of cells, such as mitochondria and the nuclear membrane (Fig. S1C&D).

### Landscape Expansion Microscopy (land-ExM)

0.0125×10^6^ cells were seeded and cultured overnight on a plasma-cleaned cover slip attached to a custom PDMS chamber, 1 mm thick and 6.5 mm in diameter, as previously described (Shi et al., 2021). The chambers are included in the LR-ExM kit (Cytoseen, cat#LR-RD488-050 or LR-MD568-050) that we purchased, and they can also be made in a lab using the SYLGARD™ 184 Silicone Elastomer Kit. To label the whole lipids, cells were fixed with 37 °C pre-warmed 3% paraformaldehyde and 0.1% Glutaraldehyde in PBS for 10 min, followed by two washes with PBS and incubation with mCLING-atto647N (Synaptic Systems, cat#710006AT1) in PBS overnight at room temperature. Since mCLING may have batch-to-batch variation, concentration optimization (1:10 to 1:100 dilution in PBS) is recommended. After mCLING staining, the cells were fixed again with 37 °C pre-warmed 3% PFA and 0.1% glutaraldehyde in PBS for 10 min, then proceeded to standard immunostaining steps if required. This included permeation with 0.1% TritonX100 in PBS (PBST), blocking with the blocking buffer (3% BSA in PBST), and primary antibody incubation overnight at 4 °C. Secondary antibodies conjugated with LR-ExM DIG trifunctional probe NHS-Digoxigenin-MA (Cytoseen, cat#LR-RD488-050 or cat#LR-MD568-050) were used to fulfill multi-color land-ExM imaging. Incubate the cells with secondary antibodies (1:50 dilution in blocking buffer) for 1h at room temperature.

To anchor and biotinylate whole proteins and anchor mCLING, incubate the cells with 2 mM NHS-Biotin-MA (Cytoseen, cat#LD-P568-050 or #LD-P488-050) in 100 mM NaHCO_3_ for 1 h, and wash with PBS 3 times. Cells are ready for gelation at this point. Cells were first incubated with pre-chilled monomer solution (8.6 g sodium acrylate, 2.5 g acrylamide, 0.15 g N,N’-methylenebisacrylamide, 11.7 g sodium chloride in 94 ml PBS buffer) on ice for 5 min then incubated with gelation solution (mixture of monomer solution, 10% (w/v) N,N,N′,N′-tetramethylethylenediamine stock solution, 10% (w/v) ammonium persulfate stock solution and water at 47:1:1:1 volume ratio) on ice for 5 min. A cover slip was applied onto the top of the PDMS chamber to seal the cell-gelation solution to avoid oxygen interruption of the gelation procedure. The cell-gelation solution was then incubated at 37 °C for 1-2 hours. Gelled cells were immersed in heat denaturation buffer (200 mM sodium dodecyl sulfate, 200 mM NaCl, and 50 mM Tris pH 6.8) for 1.5 hours at 78°C and washed with excess of water for 30 min. To fluorescently label the whole proteins, immerse the gelled cells in post-staining buffer (10 mM HEPES, 150 mM NaCl, pH 7.5) twice, 30 min each time, then incubate the gelled cells with 4 µg/mL streptavidin dyes (included in Cytoseen, cat#LD-P568-050/LD-P488-050, or JacksonimmunoResearch, cat#016-540-084) in post-staining buffer overnight. After 4 hours of washing and expansion with an excess amount of DNase/RNase-free water, the gelled cells are ready for imaging.

To compare land-ExM with other protein contextual imaging methods, after permeation or antibody staining, cells were incubated in 25 mM NHS-MA (Sigma, cat# 730300) in 100 mM sodium bicarbonate for 1 hour at room temperature, followed by three washes in PBS. After gelation and heat denaturation, gelled cells were washed with an excess amount of PBS, then incubated with 20 µg/mL NHS-Alexa Fluor 488 (Invitrogen, cat#A20000) in PBS or 1x SYPRO Orange (Invitrogen, cat# S6651) in PBS overnight. Details of reagents used can be found in the Key Resources Table in the Methods section.

For users focusing on lipid context but with a low requirement for protein imaging, glycidyl methacrylate (GMA) (Sigma, cat# 151238) can serve as an alternative to NHS-biotin-MA (Fig. 1A). Compared with the NHS-biotin-MA, using GMA as the anchor in the land-ExM workflow can result in equally high signal-to-noise ratio in lipid imaging but significantly dimmer signals in the protein channel. After permeation or antibody staining, cells were incubated with 0.04% (w/v) GMA in 100 mM sodium bicarbonate for 3 hours at room temperature, followed by washing with PBS for 3 times. After gelation and heat denaturation, gelled cells were washed with an excess amount of PBS, then incubated with 20 µg/mL NHS-Alexa Fluor 488 (Invitrogen, cat#A20000) in PBS or 1x SYPRO Orange (Invitrogen, cat# S6651) in PBS overnight.

For users focusing on protein imaging only, the heat denaturation step can be replaced by proteinase K digestion. After gelation, the gelled cell was immersed overnight in digestion buffer (50 mM Tris, 1mM EDTA, 0.5% (v/v) Triton X-100, 1M sodium chloride, pH 8) with freshly added 8 units/mL proteinase K (Sigma, cat# P4850-5ML), then washed with an excess amount of water for 30 min, excess amount of post-staining buffer twice, 30 min each time. 4 µg/mL streptavidin dye in post-staining buffer was later added and incubated with the gelled cells overnight. After expansion with an excess amount of water, the gelled cell was ready for imaging.

### land-TREx

After mCLING and antibody staining as described in land-ExM, cells were incubated with 2 mM NHS-Biotin-MA (Cytoseen, cat#LD-P568-050 or #LD-P488-050) in 100 mM NaHCO_3_ for 1 h, and then washed three times with PBS. Cells were then incubated with pre-chilled TREx monomer solution (10.3 g sodium acrylate, 14.2 g acrylamide, 0.09 g N,N’-methylenebisacrylamide in 95 ml PBS buffer) on ice for 5 min, and gelation solution (mixture of TREx monomer solution, 10% (w/v) N,N,N′,N′-tetramethylethylenediamine stock solution, 10% (w/v) ammonium persulfate stock solution and water at 64:1:1:1 volume ratio) on ice for another 5 min. A cover slip was required to place onto the top of the PDMS chamber to seal the cell-gelation solution to avoid oxygen interruption of the gelation procedure. After 2-hour gelation at 37°C in a humidity chamber, the gelled cells were immersed in heat denaturation buffer for 1.5 hours at 78°C. The gelled cell was then washed with an excess amount of water for 30 min, an excess amount of post-staining buffer for 30 min, and an excess amount of post-staining buffer for another 30 min. 4 µg/mL streptavidin dye (Cytoseen, cat#LD-P568-050 or #LD-P488-050) in post-staining buffer was added later and incubated with gelled cells overnight. After another overnight gelled cell washing with an excess amount of water, the gelled cell was expanded around 7-fold and ready for imaging.

### land-pan-ExM

0.0125×10^^6^ cells were seeded and cultured overnight on a plasma-cleaned cover slip attached to a custom PDMS chamber (0.2 mm thickness and a 6.5 mm-diameter culture area). After fixation with 3% formaldehyde and 0.1% glutaraldehyde in PBS for 15 min at room temperature and rinsing with 1xPBS once, cells were incubated with mCLING-Atto647N (Synaptic Systems, cat#710006AT1) overnight in PBS at room temperature. After mCLING staining, cells were fixed again with 3% formaldehyde and 0.1% glutaraldehyde in PBS for 15 min at room temperature. After washing with 1xPBS twice, cells were incubated with 100 mM NaHCO_3_ for 5 min, then incubated with 2 mM NHS-Biotin-MA (Cytoseen, cat#LD-P568-050 or #LD-P488-050) in 100 mM NaHCO_3_ for 1 h at room temperature, and washed with PBS 3 times. Cells were then incubated with 0.7% formaldehyde and 1% (w/v) acrylamide in PBS at 37°C for 6-7 hours, followed by PBS washing twice, 10 min each time. The cells were ready to its 1^st^ round of gelation at this point.

To fulfill the 1^st^ round of gelation, cells were first incubated with iExM 1^st^ gelation solution (19% (w/v) sodium acrylate, 10% (w/v) acrylamide, 0.1% (w/v) N,N′-(1,2-Dihydroxyethylene)bisacrylamide, 0.25% (v/v) N,N,N′,N′-tetramethylethylenediamine, and 0.25% (w/v) ammonium persulfate in PBS) for 15 min at room temperature with a coverslip sealed at the top of the PDMS chamber, then moved to a humidity chamber and incubated for 1.5 hours at 37°C. After gelation, the gelled cell was detached from the PDMS chamber and immersed in heat-denaturation buffer for 15 minutes at room temperature, then incubated for 1 hour at 78°C. Expand the gelled cells overnight with an excess amount of water. The gelled cell was ready for its 2^nd^ round of gelation at this step.

The expanded gelled cell was first incubated and gently shake with iExM 2^nd^ gelation solution (10% (w/v) acrylamide, 0.05% (w/v) N,N′-(1,2-Dihydroxyethylene)bisacrylamide, 0.05% (v/v) N,N,N′,N′-tetramethylethylenediamine, and 0.05% (w/v) ammonium persulfate in PBS) for three times, 20 min each time at room temperature, then sandwiched with two coverslips. To remove residual gelation solution, the sandwiched gelled cell was gently pressed with a Kimwipe inserted between the two coverslips. The sandwiched gelled cell was later incubated in a nitrogen-filled chamber at 37°C for 1.5 hours. After detaching the two coverslips from the gelled cell, the gelled cell was incubated with 0.7% formaldehyde and 1% (w/v) acrylamide in PBS at 37°C for 6-9 hours, followed by PBS washing three times, 30 min each time. The gelled cells were ready for their 3^rd^ round of gelation at this point.

The gelled cell was first incubated and gently shake with iExM 3^rd^ gelation solution (19% (w/v) sodium acrylate, 10% (w/v) acrylamide, 0.1% (w/v) N,N’-methylenebisacrylamide, 0.05% (v/v) N,N,N′,N′-tetramethylethylenediamine, and 0.05% (w/v) ammonium persulfate in PBS) for 4 times, 15 min each time, then sandwiched again with two coverslips. After the residual gelation solution was removed, the sandwiched gelled cell was incubated in a nitrogen-filled chamber at 37°C for 2 hours. After detaching the two coverslips from the gelled cell, the gelled cell was incubated and shaken with 0.2M NaOH for 1 hour at room temperature, followed by PBS washing 3 times, 30 minutes each time. The gelled cell was then incubated overnight with 4 µg/mL streptavidin dye (Cytoseen, cat#LD-P568-050 or #LD-P488-050) in post-staining buffer. After overnight expansion with an excess amount of water, the cell-gel was expanded to around 12-fold and was ready for imaging.

### Imaging

Gelled cells imaging was all performed on an Airyscan confocal microscope (ZEISS LSM 980 with Airyscan 2) with a 63x water-immersion objective (Zeiss LD C-Apochromat 63x/1.2 W Corr M27) with effective lateral resolution at 138 nm (measured by TetraSpeck™ Microspheres, 0.1 µm, fluorescent blue/green/orange/dark red). The effective resolution of extended samples is calculated as this microscope resolution divided by the expansion factor, ranging from 3.7 to 12.0. The highest effective resolution we could achieve was 12 nm. Airyscan SR4Y-best signal mode with 0.2 AU pinhole and 1.25 AU total detection area was used for the 3D imaging of all the samples.

### Image Rendering

Images in Fig. 4E, J, K, P, and Q were processed with a custom-written MATLAB script. Briefly, stack images were denoised by a Wiener filter, smoothed by a Gaussian filter, and then binarized. Binarized stack images were reconstructed in Imaris 10.2 using the pre-expansion dimensions. Surfaces were created, and transparency was adjusted for visualization of inner structures. The MATLAB codes are available at Xiaoyu Shi Lab’s GitHub page: https://github.com/XYShi-Lab/LandExM.git. All the other images were simply contrasted or projected using basic functions of Fiji(ImagJ).

### Quantification and statistical analysis

All images were processed and analyzed using ImageJ and Custom MATLAB code. All graphs were generated using Prism 10 (GraphPad software).

### Reagent availability

All reagents used in this work are commercially available, including land-ExM protein label, NHS-biotin-MA, and LR-ExM trifunctional antibodies. See Key Resources Table.

## Data and code availability

The MATLAB codes for 3D rendering are available at Xiaoyu Shi Lab’s GitHub page: https://github.com/XYShi-Lab/LandExM.git. All the other data analyses were performed using the open-source software Fiji (ImageJ) and are available in the published article and online supplemental material.

Fig. S1 to Fig. S7 can be found in the **Supplemental information**.

Fig. S1. mCLING optimization for lipid staining of cells.

Fig. S2. Alternative land-ExM workflow to avoid crosstalk between NHS-Biotin-MA and mCLING.

Fig. S3. land-ExM using proteinase K digestion.

Fig. S4. Land-ExM coupled with immunostaining LR-ExM for lipid vesicle identification.

Fig. S5. land-ExM coupled with immunostaining LR-ExM for membrane-bound organelles visualization.

Fig. S6. Land-ExM coupled with immunostaining for cytoskeleton visualization.

Fig. S7. Land-ExM reveals stress granules at different locations of cells.

## Acknowledgement

We thank Joerg Bewersdorf’s lab for sharing the pan-ExM protocol and Paul Tillberg’s lab for consulting on TREx. We also thank Dr. Kiryl Piatkevich for recommending Sypro Orange dye. Y.Z. and X.S. are supported by the NIH Director’s New Innovator Award (DP2GM150017, 3DP2GM150017-01S1) and the NSF Faculty Early Career Development Program (CAREER) Award (2341058). Z.Z. and Z.D. are supported by the Chan Zuckerberg Initiative (CZI) Advancing Imaging Through Collaborative Projects Award.

## Author contributions

Y. Zhuang and X. Shi conceived the study and wrote the manuscript. Y. Zhuang, Z. Zhao, Z. Dai, and X. Shi developed the methodology. X. Shi supervised the work.

## Reference

Balzarotti, F., Y. Eilers, K.C. Gwosch, A.H. Gynna, V. Westphal, F.D. Stefani, J. Elf, and S.W. Hell. 2017. Nanometer resolution imaging and tracking of fluorescent molecules with minimal photon fluxes. Science. 355:606–612.

Betzig, E., G.H. Patterson, R. Sougrat, O.W. Lindwasser, S. Olenych, J.S. Bonifacino, M.W. Davidson, J. Lippincott-Schwartz, and H.F. Hess. 2006. Imaging intracellular fluorescent proteins at nanometer resolution. Science. 313:1642–1645.

Bianchini, P., L. Pesce, and A. Diaspro. 2021. Expansion microscopy at the nanoscale: The nuclear pore complex as a fiducial landmark. Methods Cell Biol. 161:275–295.

Bouteille, M.B.C.A., and D. Hemon. 1979. Structural Relationship Between the Nucleolus and the Nuclear-Envelope. J Ultrastruct Res. 68:328–340.

Bussi, C., A. Mangiarotti, C. Vanhille-Campos, B. Aylan, E. Pellegrino, N. Athanasiadi, A. Fearns, A. Rodgers, T. Franzmann, A. Saric, R. Dimova, and M. Gutierrez. 2023. Stress granules plug and stabilize damaged endolysosomal membranes. Nature. 623:1062–1069.

Chang, J.-B., F. Chen, Y.-G. Yoon, E.E. Jung, H. Babcock, J.S. Kang, S. Asano, H.-J. Suk, N. Pak, P.W. Tillberg, A.T. Wassie, D. Cai, and E.S. Boyden. 2017. Iterative expansion microscopy. Nature Methods. 14:593–599.

Chen, F., P.W. Tillberg, and E.S. Boyden. 2015. Expansion microscopy. Science. 347:543–548.

Chen, F., A.T. Wassie, A.J. Cote, A. Sinha, S. Alon, S. Asano, E.R. Daugharthy, J.B. Chang, A. Marblestone, G.M. Church, A. Raj, and E.S. Boyden. 2016. Nanoscale imaging of RNA with expansion microscopy. Nat Methods. 13:679–684.

Chozinski, T.J., A.R. Halpern, H. Okawa, H.-J. Kim, G.J. Tremel, R.O.L. Wong, and J.C. Vaughan. 2016. Expansion microscopy with conventional antibodies and fluorescent proteins. Nature Methods. 13:485–488.

Cui, Y., G. Yang, D.R. Goodwin, C.H. O’Flanagan, A. Sinha, C. Zhang, K.E. Kitko, T.W. Shin, D. Park, S. Aparicio, C.I.G.C. Consortium, and E.S. Boyden. 2023. Expansion microscopy using a single anchor molecule for high-yield multiplexed imaging of proteins and RNAs. PLOS ONE. 18:e0291506.

Damstra, H.G.J., B. Mohar, M. Eddison, A. Akhmanova, L.C. Kapitein, and P.W. Tillberg. 2022. Visualizing cellular and tissue ultrastructure using Ten-fold Robust Expansion Microscopy (TREx). Elife. 11:e73775.

Damstra, H.G.J., J.B. Passmore, A.K. Serweta, I. Koutlas, M. Burute, F.J. Meye, A. Akhmanova, and L.C. Kapitein. 2023. GelMap: intrinsic calibration and deformation mapping for expansion microscopy. Nat Methods. 20:1573–1580.

de Boer, P., J.P. Hoogenboom, and B.N. Giepmans. 2015. Correlated light and electron microscopy: ultrastructure lights up! Nat Methods. 12:503–513.

Godman, G.C., C. Morgan, P.M. Breitenfeld, and H.M. Rose. 1960. A correlative study by electron and light microscopy of the development of type 5 adenovirus. II. Light microscopy. J Exp Med. 112:383–402.

Grunewald, K., P. Desai, D.C. Winkler, J.B. Heymann, D.M. Belnap, W. Baumeister, and A.C. Steven. 2003. Three-dimensional structure of herpes simplex virus from cryo-electron tomography. Science. 302:1396–1398.

Gustafsson, M.G.L. 2000. Surpassing the lateral resolution limit by a factor of two using structured illumination microscopy. *J*. Microscopy. 198:82–87.

Hauser, M., M. Wojcik, D. Kim, M. Mahmoudi, W. Li, and K. Xu. 2017. Correlative Super-Resolution Microscopy: New Dimensions and New Opportunities. Chem Rev. 117:7428–7456.

Hell, S.W., and J. Wichmann. 1994. Breaking the diffraction resolution limit by stimulated emission: stimulated-emission-depletion fluorescence microscopy. Opt Lett. 19:780–782.

Huang, B., S.A. Jones, B. Brandenburg, and X. Zhuang. 2008. Whole-cell 3D STORM reveals interactions between cellular structures with nanometer-scale resolution. Nat Methods. 5:1047–1052.

Karagiannis, E.D., J.S. Kang, T.W. Shin, A. Emenari, S. Asano, L. Lin, E.K. Costa, I.G.C. Consortium, A.H. Marblestone, N. Kasthuri, and E.S. Boyden. 2019. Expansion Microscopy of Lipid Membranes. bioRxiv:829903.

King, M.C., C.P. Lusk, and N.R. Ader. 2025. Sense, plug, and seal: proteins as both rapid responders and constitutive barriers supporting organelle compartmentalization. Molecular Biology of the Cell. 36:pe6.

Klimas, A., B.R. Gallagher, P. Wijesekara, S. Fekir, E.F. DiBernardo, Z. Cheng, D.B. Stolz, F. Cambi, S.C. Watkins, S.L. Brody, A. Horani, A.L. Barth, C.I. Moore, X. Ren, and Y. Zhao. 2023. Magnify is a universal molecular anchoring strategy for expansion microscopy. Nature Biotechnology:1–12.

Ku, T., J. Swaney, J.-Y. Park, A. Albanese, E. Murray, J.H. Cho, Y.-G. Park, V. Mangena, J. Chen, and K. Chung. 2016. Multiplexed and scalable super-resolution imaging of three-dimensional protein localization in size-adjustable tissues. Nature Biotechnology. 34:973–981.

Laporte, M.H., N. Klena, V. Hamel, and P. Guichard. 2022. Visualizing the native cellular organization by coupling cryofixation with expansion microscopy (Cryo-ExM). Nat Methods. 19:216–222.

Li, H., A.R. Warden, J. He, G. Shen, and X. Ding. 2022. Expansion microscopy with ninefold swelling (NIFS) hydrogel permits cellular ultrastructure imaging on conventional microscope. Science Advances. 8:eabm4006.

Liu, S., X. Zhang, X. Yao, G. Wang, S. Huang, P. Chen, M. Tang, J. Cai, Z. Wu, Y. Zhang, R. Xu, K. Liu, K. He, Y. Wang, L. Jiang, Q.A. Wang, L. Rui, J. Liu, and Y. Liu. 2024. Mammalian IRE1alpha dynamically and functionally coalesces with stress granules. Nat Cell Biol. 26:917–931.

M’Saad, O., and J. Bewersdorf. 2020. Light microscopy of proteins in their ultrastructural context. Nat Commun. 11:3850.

Malhas, A., C. Goulbourne, and D.J. Vaux. 2011. The nucleoplasmic reticulum: form and function. Trends in Cell Biology. 21:362–373.

Mao, C., M.Y. Lee, J.-R. Jhan, A.R. Halpern, M.A. Woodworth, A.K. Glaser, T.J. Chozinski, L. Shin, J.W. Pippin, S.J. Shankland, J.T.C. Liu, and J.C. Vaughan. 2020. Feature-rich covalent stains for super-resolution and cleared tissue fluorescence microscopy. Science Advances. 6:eaba4542.

M’Saad, O., and J. Bewersdorf. 2020. Light microscopy of proteins in their ultrastructural context. Nature Communications. 11:3850.

Narayan, K., and S. Subramaniam. 2015. Focused ion beams in biology. Nat Methods. 12:1021–1031.

Nicchitta, C.V. 2024. An emerging role for the endoplasmic reticulum in stress granule biogenesis. Semin Cell Dev Biol. 156:160–166.

Obara, C.J., J. Nixon-Abell, A.S. Moore, F. Riccio, D.P. Hoffman, G. Shtengel, C.S. Xu, K. Schaefer, H.A. Pasolli, J.B. Masson, H.F. Hess, C.P. Calderon, C. Blackstone, and J. Lippincott-Schwartz. 2024. Motion of VAPB molecules reveals ER-mitochondria contact site subdomains. Nature. 626:169–176.

Pavani, S.R., M.A. Thompson, J.S. Biteen, S.J. Lord, N. Liu, R.J. Twieg, R. Piestun, and W.E. Moerner. 2009. Three-dimensional, single-molecule fluorescence imaging beyond the diffraction limit by using a double-helix point spread function. Proc Natl Acad Sci U S A. 106:2995–2999.

Pincus, D., and S.A. Oakes. 2024. Unfolding emergency calls stress granules to the ER. Nat Cell Biol. 26:845–846.

Pownall, M.E., L. Miao, C.E. Vejnar, O. M’Saad, A. Sherrard, M.A. Frederick, M.D.J. Benitez, C.W. Boswell, K.S. Zaret, J. Bewersdorf, and A.J. Giraldez. 2023. Chromatin expansion microscopy reveals nanoscale organization of transcription and chromatin. Science. 381:92–100.

Raimondi, A., N. Ilacqua, and L. Pellegrini. 2023. Liver inter-organelle membrane contact sites revealed by serial section electron tomography. Methods in Cell Biology.177:101-123.

Revelo, N.H., D. Kamin, S. Truckenbrodt, A.B. Wong, K. Reuter-Jessen, E. Reisinger, T. Moser, and S.O. Rizzoli. 2014. A new probe for super-resolution imaging of membranes elucidates trafficking pathways. J Cell Biol. 205:591–606.

Rigort, A., F.J. Bauerlein, E. Villa, M. Eibauer, T. Laugks, W. Baumeister, and J.M. Plitzko. 2012. Focused ion beam micromachining of eukaryotic cells for cryoelectron tomography. Proc Natl Acad Sci U S A. 109:4449–4454.

Rust, M.J., M. Bates, and X. Zhuang. 2006. Sub-diffraction-limit imaging by stochastic optical reconstruction microscopy (STORM). Nat Methods. 3:793–795.

Scheckenbach, M., J. Bauer, J. Zähringer, F. Selbach, and P. Tinnefeld. 2020. DNA origami nanorulers and emerging reference structures. APL Materials. 8:110902.

Schmidt, R., T. Weihs, C.A. Wurm, I. Jansen, J. Rehman, S.J. Sahl, and S.W. Hell. 2021. MINFLUX nanometer-scale 3D imaging and microsecond-range tracking on a common fluorescence microscope. Nat Commun. 12:1478.

Scorrano, L., M. De Matteis, S. Emr, F. Giordano, G. Hajnóczky, B. Kornmann, L. Lackner, T. Levine, L. Pellegrini, K. Reinisch, R. Rizzuto, T. Simmen, H. Stenmark, C. Ungermann, and M. Schuldiner. 2019. Coming together to define membrane contact sites. Nature Communications. 10.

Shaib, A.H., A.A. Chouaib, R. Chowdhury, J. Altendorf, D. Mihaylov, C. Zhang, D. Krah, V. Imani, R.K.W. Spencer, S.V. Georgiev, N. Mougios, M. Monga, S. Reshetniak, T. Mimoso, H. Chen, P. Fatehbasharzad, D. Crzan, K.A. Saal, M.M. Alawieh, N. Alawar, J. Eilts, J. Kang, A. Soleimani, M. Muller, C. Pape, L. Alvarez, C. Trenkwalder, B. Mollenhauer, T.F. Outeiro, S. Koster, J. Preobraschenski, U. Becherer, T. Moser, E.S. Boyden, A.R. Aricescu, M. Sauer, F. Opazo, and S.O. Rizzoli. 2024. One-step nanoscale expansion microscopy reveals individual protein shapes. Nat Biotechnol:10.1038/s41587-41024-02431-41589.

Shi, X., Q. Li, Z. Dai, A.A. Tran, S. Feng, A.D. Ramirez, Z. Lin, X. Wang, T.T. Chow, J. Chen, D. Kumar, A.R. McColloch, J.F. Reiter, E.J. Huang, I.B. Seiple, and B. Huang. 2021. Label-retention expansion microscopy. J Cell Biol. 220:e202105067.

Shin, T., H. Wang, C. Zhang, B. An, Y. Lu, E. Zhang, X. Lu, E. Karagiannis, J. Kang, A. Emenari, P. Symvoulidis, S. Asano, L. Lin, E. Costa, A. Marblestone, N. Kasthuri, L. Tsai, and E. Boyden. 2025. Dense, continuous membrane labeling and expansion microscopy visualization of ultrastructure in tissues. Nature Communications. 16.

Sun, D.-e., X. Fan, Y. Shi, H. Zhang, Z. Huang, B. Cheng, Q. Tang, W. Li, Y. Zhu, J. Bai, W. Liu, Y. Li, X. Wang, X. Lei, and X. Chen. 2021. Click-ExM enables expansion microscopy for all biomolecules. Nature Methods. 18:107–113.

Tillberg, P.W., F. Chen, K.D. Piatkevich, Y. Zhao, C.-C. Yu, B.P. English, L. Gao, A. Martorell, H.-J. Suk, F. Yoshida, E.M. DeGennaro, D.H. Roossien, G. Gong, U. Seneviratne, S.R. Tannenbaum, R. Desimone, D. Cai, and E.S. Boyden. 2016. Protein-retention expansion microscopy of cells and tissues labeled using standard fluorescent proteins and antibodies. Nature Biotechnology. 34:987–992.

Truckenbrodt, S., M. Maidorn, D. Crzan, H. Wildhagen, S. Kabatas, and S.O. Rizzoli. 2018. X10 expansion microscopy enables 25-nm resolution on conventional microscopes. EMBO reports. 19:e45836.

Vanheusden, M., R. Vitale, R. Camacho, K.P.F. Janssen, A. Acke, S. Rocha, and J. Hofkens. 2020. Fluorescence Photobleaching as an Intrinsic Tool to Quantify the 3D Expansion Factor of Biological Samples in Expansion Microscopy. ACS Omega. 5:6792–6799.

Wang, S., T.W. Shin, H.B. Yoder, 2nd, R.B. McMillan, H. Su, Y. Liu, C. Zhang, K.S. Leung, P. Yin, L.L. Kiessling, and E.S. Boyden. 2024. Single-shot 20-fold expansion microscopy. Nat Methods. 21:2128–2134.

Wheeler, J., T. Matheny, S. Jain, R. Abrisch, and R. Parker. 2016. Distinct stages in stress granule assembly and disassembly. Elife. 5.

White, B.M., P. Kumar, A.N. Conwell, K. Wu, and J.M. Baskin. 2022. Lipid Expansion Microscopy. Journal of the American Chemical Society.

Wu, H., P. Carvalho, and G.K. Voeltz. 2018. Here, there, and everywhere: The importance of ER membrane contact sites. Science. 361:eaan5835.

Xu, C.S., K.J. Hayworth, Z. Lu, P. Grob, A.M. Hassan, J.G. Garcia-Cerdan, K.K. Niyogi, E. Nogales, R.J. Weinberg, and H.F. Hess. 2017. Enhanced FIB-SEM systems for large-volume 3D imaging. Elife. 6:e25916.

Zhuang, Y., X. Guo, O.V. Razorenova, C.E. Miles, W. Zhao, and X. Shi. 2024. Coaching ribosome biogenesis from the nuclear periphery. bioRxiv:2024.2006.2021.597078.

Zhuang, Y., and X. Shi. 2023. Expansion microscopy: A chemical approach for super-resolution microscopy. Curr Opin Struct Biol. 81:102614.

