## Supplemental Information for "Landscape Expansion Microscopy Reveals Interactions between Membrane and Phase-Separated Organelles"

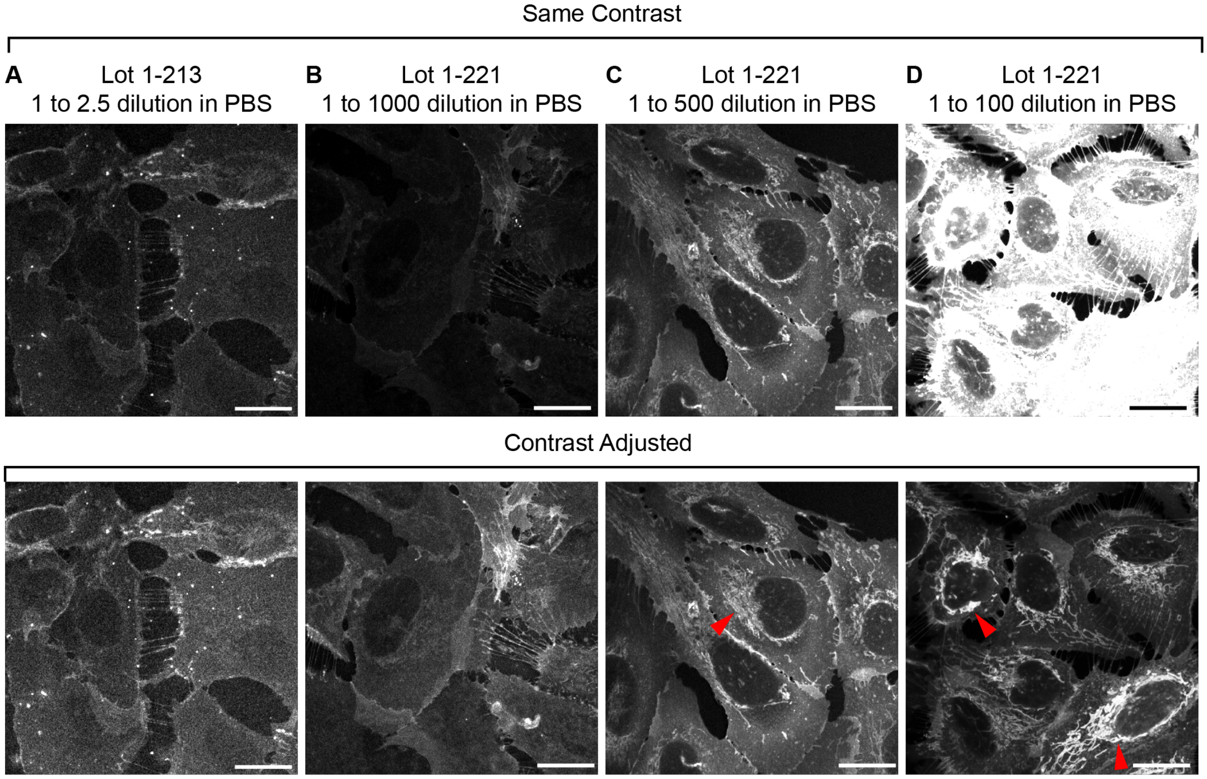


**Figure S1. mCLING optimization for lipid staining of cells.**

(A-D) Airyscan images of U2OS cells stained with different batches of mCLING at different dilution factors. Scale bars: 20 µm. Red arrowheads indicate lipid structures in the cytoplasm. All images were taken with an Airyscan microscope.

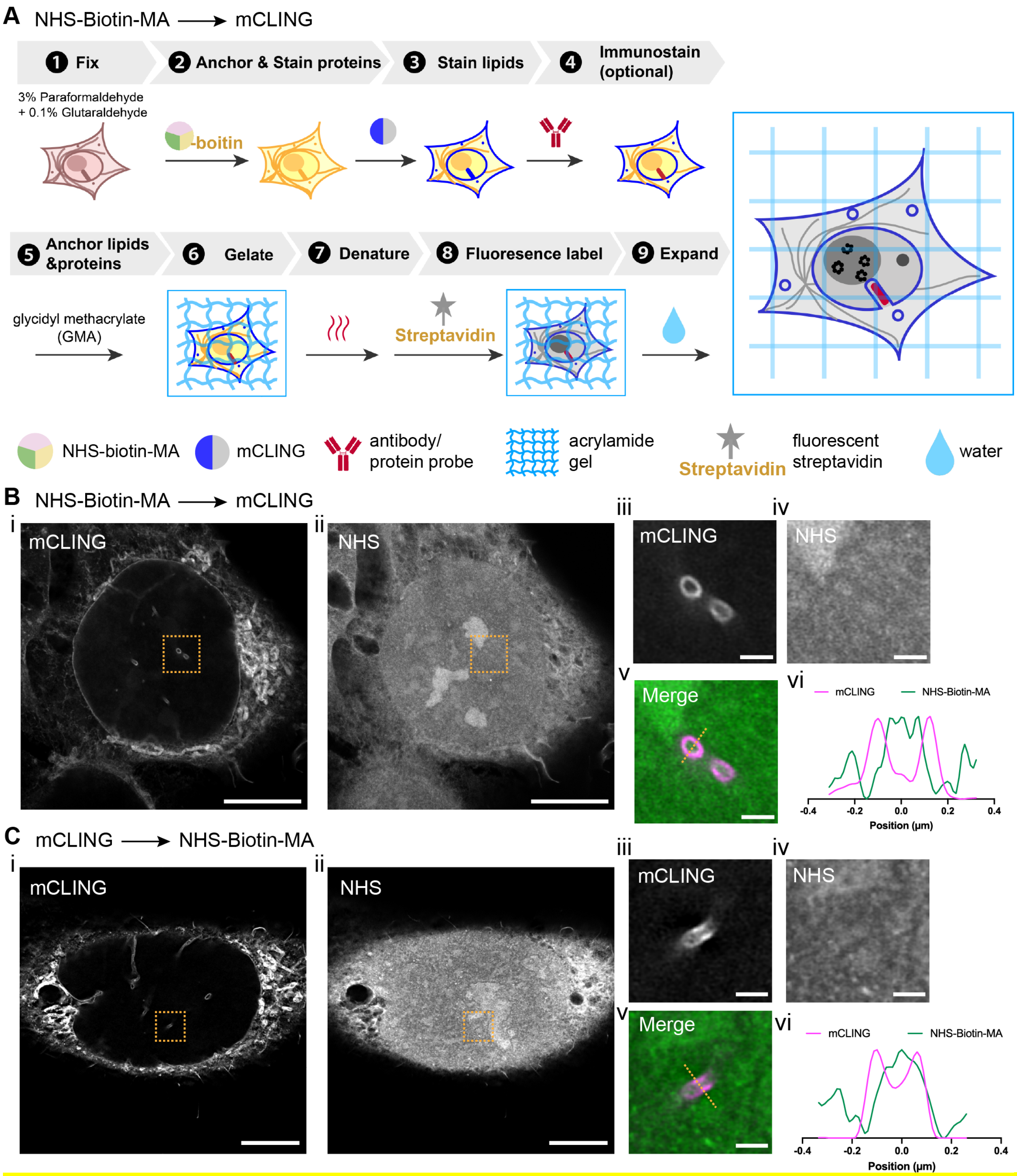


**Figure S2. Alternative land-ExM workflow to avoid crosstalk between NHS-Biotin-MA and mCLING.** (A) Alternative workflow of land-ExM. (B) i and ii: land-ExM images of U2OS cells stained first with NHS-Biotin-MA then mCLING. iii to v: Magnified images of boxes in i and ii. vi: Normalized intensity profile along the orange line in v. Scale bar: 5 µm in pre-expansion unit. Linear expansion factor: 4.0 (i and ii). 0.5 µm in pre-expansion unit. Linear expansion factor: 4..0 (iii to v). (C) i and ii: land-ExM images of U2OS cells stained first with mCLING then NHS-Biotin-MA. iii to v: Magnified images of orange boxes in i and ii. vi: Normalized intensity profile along the orange line in v. Scale bar: 5 µm in pre-expansion unit. Linear expansion factor: 4 (i and ii). 0.5 µm in pre-expansion unit. Linear expansion factor: 4.0 (iii to v). All images were taken with an Airyscan microscope.

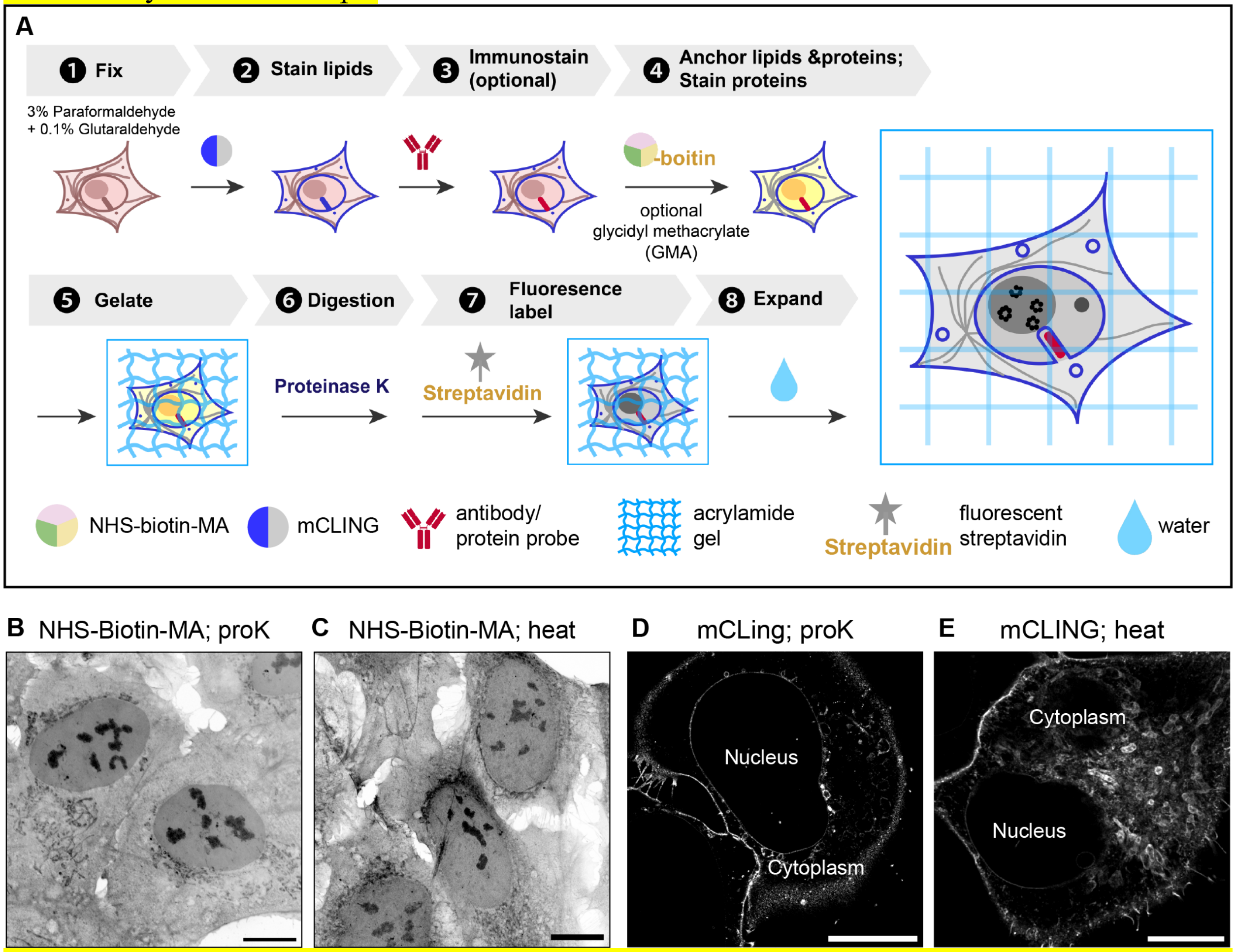


**Figure S3. land-ExM using proteinase K digestion.** (A) Workflow of land-ExM using proteinase K digestion to homogenize cells instead of heat denaturation. (B) land-ExM protein image of U2OS cells with proteinase K digestion (proK). Scale bar: 10 µm in pre-expansion unit. Linear expansion factor: 4.0. (C) land-ExM protein image of U2OS cells with heat denaturation (heat). Scale bar: 10 µm in pre-expansion unit. Linear expansion factor: 4.0. (D) land-ExM lipid image of U2OS cell with proteinase K digestion (proK). Scale bar: 10 µm in pre-expansion unit. Linear expansion factor: 4.0. (E) Land-ExM lipid image of U2OS cells with heat denaturation (heat). Scale bar: 10 µm in pre-expansion unit. Linear expansion factor: 4.0. All images were taken with an Airyscan microscope.

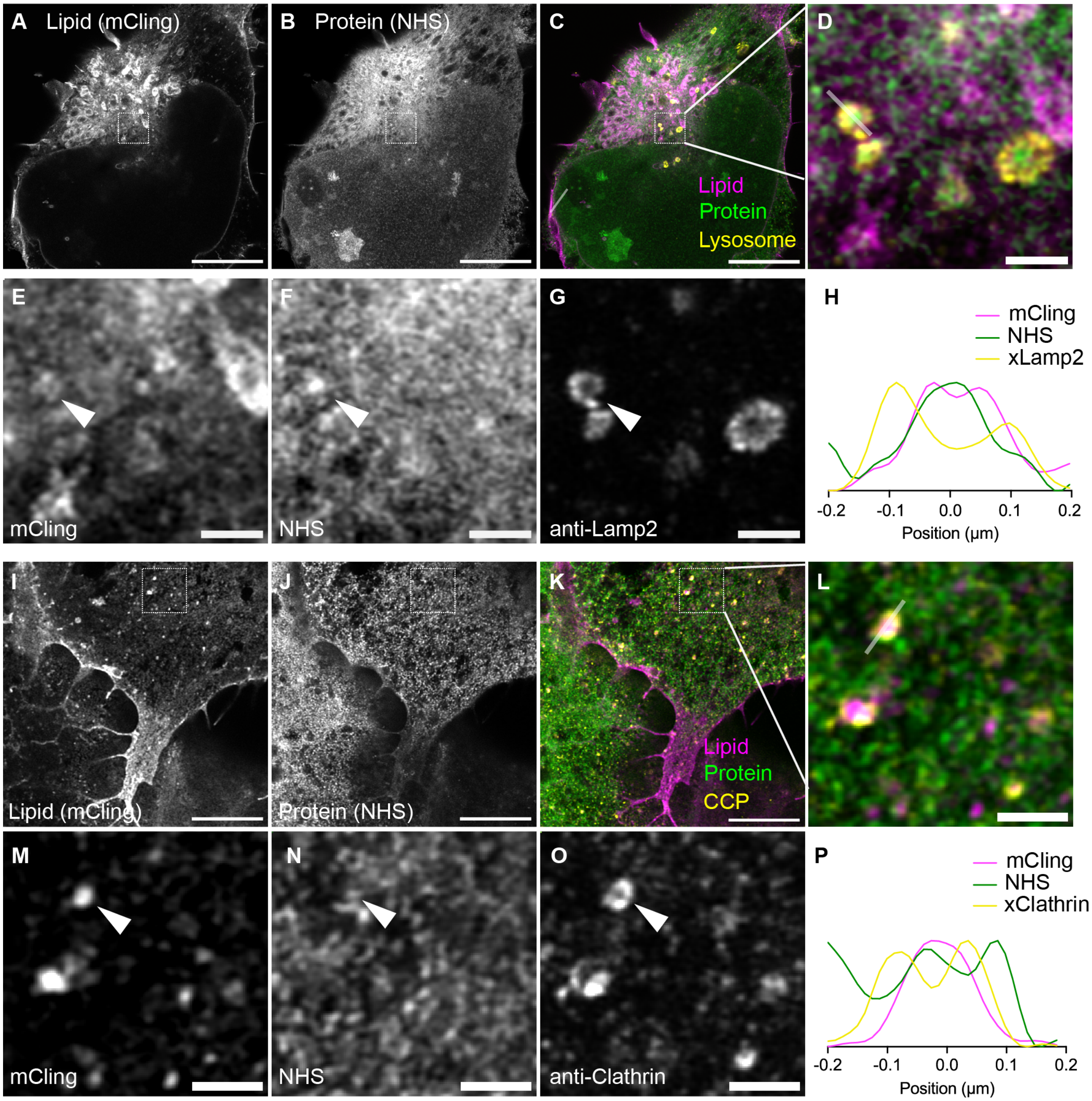


**Figure S4. Land-ExM coupled with immunostaining LR-ExM for lipid vesicle identification.**

(A-C) Land-ExM lipid (magenta) and protein (green) images of U2OS cells immunostained with anti-Lamp2 antibodies (yellow). The anti-Lamp2 antibodies are labeled LR-ExM 2nd antibodies, which are 2nd antibodies conjugated with NHS-digoxigenin-MA. Scale bar: 5 μm in pre-expansion unit. Linear expansion factor: 4.

(D-G) Magnified images of (A-C) showing details of lysosomes. Scale bar: 500 nm in pre-expansion unit.

(H) Intensity profile along the grey line across the lysosome in image (D).

(I-K) Land-ExM lipid (magenta) and protein (green) images of U2OS cells immunostained with anti-clathrin antibodies (yellow). The anti-clathrin antibodies are labeled LR-ExM 2nd antibodies,

which are 2nd antibodies conjugated with NHS-digoxigenin-MA. Scale bar: 5 µm in pre-expansion unit. Linear expansion factor: 4.
(L-O) Magnified images of (I-K) showing details of clathrin-coated pits. Scale bar: 500 nm in pre-expansion unit.
(P) Intensity profile along the grey line across the clathrin-coated pit in image (L).
All images were taken with an Airyscan microscope.

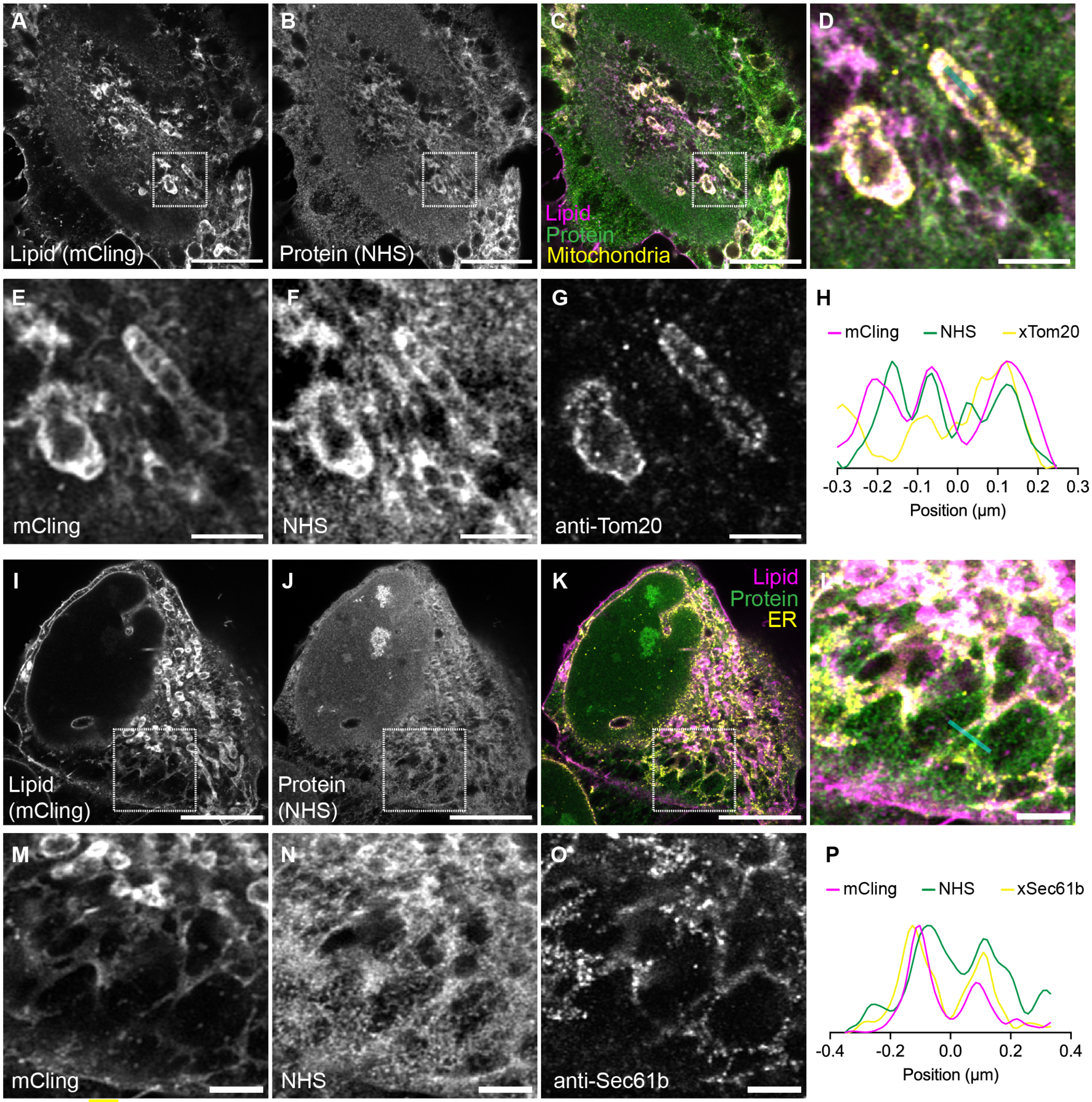


**Figure S5. land-ExM coupled with immunostaining LR-ExM for membrane-bound organelles visualization.**
(A-C) Land-ExM total lipid (magenta) and protein (green) images of U2OS cells immunostained with anti-Tom20 antibodies (yellow). The anti-Tom20 antibodies are labeled LR-ExM 2nd

antibodies, which are 2nd antibodies conjugated with NHS-digoxigenin-MA. Scale bar: 5 µm in pre-expansion unit. Linear expansion factor: 4.
(D-G) Magnified images of (A-C) showing details of mitochondria. Scale bar: 1 µm in pre-expansion unit.
(H) Intensity profile along the cyan line across the mitochondria in image (D).
(I-K) Land-ExM lipid (magenta) and protein (green) images of U2OS cells immunostained with anti-Sec61b antibodies (yellow). The anti-Sec61b antibodies are labeled LR-ExM 2nd antibodies, which are 2nd antibodies conjugated with NHS-digoxigenin-MA. Scale bar: 5 µm in pre-expansion unit. Linear expansion factor: 4.
(L-O) Magnified in images of (I-K) showing details of endoplasmic reticulum (ER). Scale bar: 1 µm in pre-expansion unit.
(P) Intensity profile along the cyan line across the ER in image (L).
All images were taken with an Airyscan microscope.

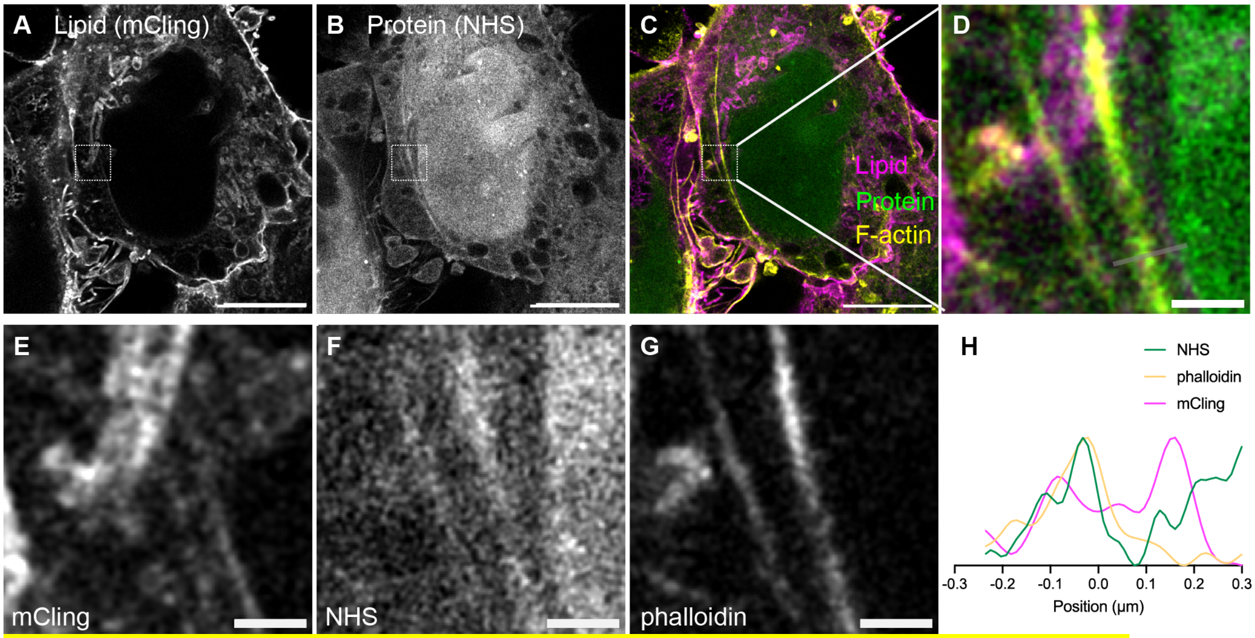


**Figure S6. Land-ExM coupled with immunostaining for cytoskeleton visualization.**
(A-C) Land-ExM total lipid (magenta) and protein (green) images of breast cancer cells stained with phalloidin-FITC and anti-FITC antibody (yellow). Scale bar: 5 µm in pre-expansion unit. Linear expansion factor: 3.8.
(D-G) Zoom in images of (A-C) showing details of F-actin. Scale bar: 500 nm in pre-expansion unit.
(H) Intensity profile along the grey line in image (D).
All images were taken with an Airyscan microscope.

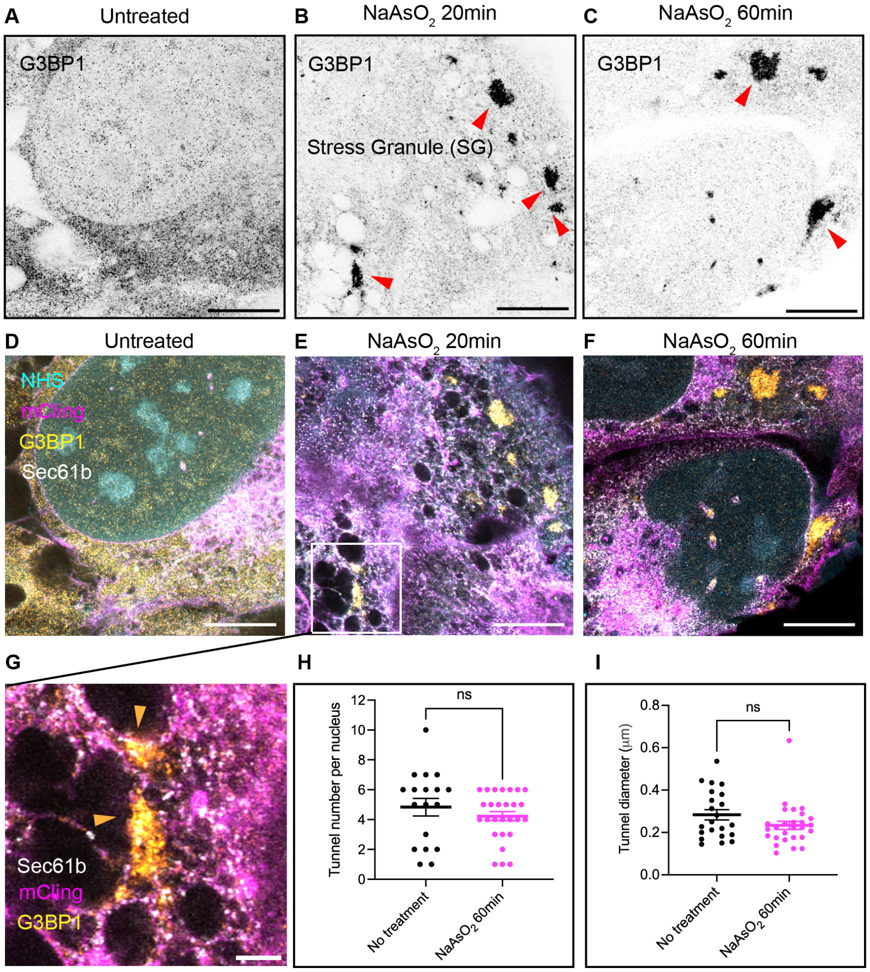


**Figure S7. Land-ExM reveals stress granules at different locations of cells.** (A-C) Land-ExM images of U2OS cells untreated or treated with NaAsO2 for 20 min or 60 min, then immunostained with anti-G3BP1 antibody. Scale bar: 5 µm in pre-expansion unit. Linear expansion factor: 4. (D-F) Land-ExM images of U2OS cells stained with mCLING (magenta), NHS ester dye (cyan), and immunostained with anti-G3BP1 (yellow) and anti-Sec61b (white) antibodies. Cells were untreated or treated with NaAsO2 for 20 min or 60 min. Scale bar: 5 µm in pre-expansion unit. Linear expansion factor: 4. (G) Magnified images of (E) showing stress granules (SG) formed adjacent to ER (orange arrowheads). Scale bar: 1 µm in pre-expansion unit.

(H) Analysis of the number of nuclear tunnels per cell with or without 60 min NaAsO2 treatment. Each bar represents the mean ± standard error of more than 18 cells. The ns indicates p>0.05 by Welch's t-test.

(I) Analysis of the diameter of nuclear tunnels in cells with or without 60 min NaAsO2 treatment. Each bar represents the mean ± standard error of more than 20 cells. ns indicates p>0.05 by Welch's t-test.

All images were taken with an Airyscan microscope.
